# Rhes, a Striatal-Enriched Protein, Promotes Mitophagy Via Nix

**DOI:** 10.1101/703934

**Authors:** Manish Sharma, Uri Nimrod Ramirez Jarquin, Oscar Rivera, Melissa Karantzis, Mehdi Eshraghi, Neelam Shahani, Vishakha Sharma, Ricardo Tapia, Srinivasa Subramaniam

**Affiliations:** The Scripps Research Institute, Department of Neuroscience, Jupiter, FL 33458, USA; The Scripps Research Institute, Metabolic Core, Jupiter, FL 33458, USA; División de Neurociencias, Instituto de Fisiología Celular, Universidad Nacional Autónoma de México, Ciudad de México-04510, México

**Author notes:** equal contributions (Manish Sharma, Uri Nimrod Ramirez Jarquin).

**Keywords:** Striatal-neuronal vulnerability, SDHA, mitophagy-ligand, SUMO-E3 ligase, lysosomes, live-cell imaging, electron microscopy, mitophagosomes, cycloheximide, protein-protein interaction, tunneling nanotubes.

## Abstract

Elimination of dysfunctional mitochondria via mitophagy is essential for cell survival and neuronal functions. But, how impaired mitophagy participates in tissue-specific vulnerability in the brain remains unclear. Here we discovered that Rhes, a striatal-enriched protein, is a major regulator of mitophagy in the striatum. Rhes predominantly interact with dysfunctional mitochondria and degrades them via mitophagy, and this function is exacerbated by the striatal toxin, 3-nitropropionic acid (3-NP). 3-NP induces mitochondrial swelling, loss of cristae and neuronal cell death only in WT but not Rhes KO striatum. Mechanistically, Rhes disrupts the mitochondrial membrane potential (ΔΨ*_m_*) and interacts with mitophagy receptor, Nix. In Nix KO cells, Rhes fails to disrupt ΔΨ*_m_* or eliminate dysfunctional mitochondria. Moreover, Rhes travels to the neighboring cell and associates with dysfunctional mitochondria via Nix. Collectively, Rhes is a major regulator of mitophagy via Nix which may determine striatal vulnerability in the brain.

## Introduction

Understanding mitophagy mechanisms and its dysregulation leading to pathological abnormalities in human diseases is a major challenge in modern biology. Reduced mitochondrial functions are linked to aging and many neurodegenerative disorders. Parkinson disease (PD) is the best example linked to mitochondrial dysfunction, because the most vulnerable neurons of PD, the substantia nigra pars compacta, show mitochondrial abnormalities, mitochondrial toxins like 1-methyl-4-phenyl-1,2,3,6-tetrahydropyridine (MPTP) elicits PD-like neurodegeneration and symptoms, and familial PD genes, such as PTEN-induced kinase-1 (PINK1), Parkin, and leucine rich repeat kinase 2 (LRRK2), which are implicated in mitochondrial dysfunction (Barodia et al., 2017; Vives-Bauza et al., 2010; Wang et al., 2012). Decline in mitochondrial enzyme activity is reported more in Alzheimer disease (AD) and amyotrophic lateral sclerosis (ALS) patients compared to control subjects (Zhang et al., 2016). Impaired mitochondrial function is observed in Huntington disease (HD) as patients continue to drop weight, even though they have a high caloric intake (Jenkins et al., 1993). Oligomers of amyloid, mutant Superoxide Dismutase 1 (SOD1), and mutant Huntingtin (mHTT) can affect mitochondrial membrane potentials and impair mitochondria trafficking and function (Browne et al., 1997; Mattiazzi et al., 2002; Oliveira, 2010; Panov et al., 2002; Shi et al., 2010; Yan et al., 1997; Zhang et al., 2018). Despite these important studies, how mitochondrial dysfunction leads to selective neuronal vulnerability remains unknown. For example, although, the familial PINK1 and Parkin mutants are ubiquitously expressed (Truban et al., 2017), the mechanisms by which they elicit lesion in the substantia nigra remains largely unclear. Similarly, how mitochondrial toxins induce tissue-specific lesion in the brain is also unclear. It is well known that 3 nitropropionic acid (3-NP) promotes lesion in the striatum but not in the cortex or cerebellar neurons, resulting in HD-like motor deficits (Brouillet et al., 1995; Brouillet et al., 2005). Similarly, MPTP or rotenone promotes substantia nigra pars compacta neurodegeneration, sparing the cortex or striatum, resulting in PD-like symptoms(Cannon et al., 2009; Langston et al., 1983; Parker et al., 1989). The major difference between these toxins that promote brain region-specific lesion are that 3-NP blocks complex II, Succinate dehydrogenase (SDH) (Huang et al., 2006), whereas rotenone and MPTP block complex I (NADH:ubiquinone oxidoreductase) (Nicklas et al., 1985; Ramsay et al., 1991). Strikingly, complex II is the only oxidative complex of the respiratory chain that does not add to the proton gradient. Thus, what determines the substantia nigra neuron to MPTP-induced lesion and striatal neuron to 3-NP induced lesion remains unclear. The differences in their brain penetrability may not account for the tissue-specific lesion as these toxins block mitochondria throughout the brain and peripheral tissue. Therefore, although mitochondrial dysfunction is ubiquitous, it is insufficient to elicit selective neuronal death, suggesting that there are additional mechanisms that may play role in the brain (Banerjee et al., 2009; Thomas and Mohanakumar, 2004).

Rhes belongs to small GTPase family of proteins highly enriched in the brain’s striatum, which controls psychiatric, cognitive and motor functions. Rhes is induced by thyroid hormones and can inhibit the cAMP/PKA pathway, dopaminergic signaling, and N-type Ca^2+^ channels (Cav 2.2)(Errico et al., 2008; Ghiglieri et al., 2015; Harrison and He, 2011; Vargiu et al., 2004). Over the years, we have found several new roles for Rhes in the striatum. Rhes can regulate the mammalian target of rapamycin complex 1 (mTORC1), SUMOylation and HD toxicity in cell and mouse models (Subramaniam et al., 2010; Subramaniam et al., 2012; Subramaniam et al., 2009; Subramaniam and Snyder, 2011; Swarnkar et al., 2015). Independent studies show a toxic link for Rhes in striatal toxicity in various models of HD (Argenti, 2014; Baiamonte et al., 2013; Lu and Palacino, 2013; Okamoto et al., 2009; Sbodio et al., 2013; Seredenina et al., 2011; Subramaniam et al., 2009). Rhes KO mice are also resistant to 3-NP induced striatal lesion (Mealer et al., 2013). But the mechanisms by which Rhes promotes striatal vulnerability in the brain remains unclear.

Here we report that Rhes is a critical regulator of mitophagy in the striatum. Using ultrastructure, biochemical, and cell and molecular biology tools, we demonstrate Rhes upregulates mitophagy via Nix receptor leading to striatal cell death. Our study reveals novel mitophagy mechanisms by which Rhes promotes striatal vulnerability in the brain.

## Results

### Rhes associates with mitochondria and lysosomes in vivo and in vitro

We took an unbiased approach by comparing our previously identified in vivo striatal interactome of Rhes (Shahani et al., 2016) with a publicly available database of mitochondrial (www.mrc-mbu.cam.ac.uk/impi) and lysosomal proteins (Wyant et al., 2018). We found 64 (3.5%) and 20 (1.8%), out of 307 high confidence striatal interactor of Rhes with potential association to mitochondria (example, Vdac1, Rhot1) and lysosomes (Atp6v0d1, Nid1), respectively (Fig. 1A, B). Next, using sucrose gradient density, we separated organelles from the striatal brain tissue homogenate from WT and Rhes KO and found that endogenous Rhes co-sedimented in mitochondrial [as detected by Succinate dehydrogenase subunit A (SDHA)] and lysosomal [(Microtubule associated protein light chain 3 (LC3) and lysosome associated membrane protein 1 (LAMP1)] fractions in WT striatum, and as expected no Rhes signal was detected in KO (Fig. 1C). In order to further understand the Rhes’ mitochondrial role, we employed striatal neuronal cell lines (a. k. a. STR*^Hdh^*) isolated from the striatum of knock-in mice, which contain a targeted insertion of a chimeric human–mouse Htt exon 1 with 7 CAG (control) (Trettel et al., 2000). These cells do not express endogenous Rhes, so we think it is a suitable model for studying the function of exogenously added Rhes, as it is the only source of Rhes, and it would not compete with endogenous function of Rhes. Similar to endogenous Rhes in the striatum, the cultured in striatal neuronal cell line expressing GFP-Rhes, not GFP alone (control), also co-sedimented in the mitochondrial and lysosomal fractions (Fig. 1D). Next, we carried live-cell time-lapse fluorescence confocal microscopy to determine Rhes interaction and function with mitochondria and lysosomes. We found that: a) Rhes interacted preferentially with globular mitochondria in primary striatal neurons processes (Fig. 1E, insets e1, e2, and deconvoluted images, e1-3D, e2-3D) and striatal neuronal cell line cytoplasm (Fig. 1F, insets f1, f2 and f2-3D) but not with the elongated mitochondria (arrowheads, Fig. 1E, F); b) Rhes formed a circular structure around the globular mitochondria (arrow, f2-3D), strongly colocalizes with it (Fig. 1F); c) Rhes colocalizes with lysosomes in primary striatal neuronal processes (Fig. 1h, arrow, h1-3D) as well as striatal neuronal cell lines (Fig. 1I, arrow, i1-3D, 1J); and d) triple staining reveals Rhes localizes with both mitochondria and lysosomes in striatal neuronal cell (Fig. 1K, insets k1 and k1-3D). GFP alone did not show colocalization with mitotracker or lysosome in striatal neuronal cells or primary neurons (Fig. S1). Next, we performed live-cell imaging of striatal neuronal cells transfected with GFP-Rhes (green) and co-stained with mitotracker (red) and lysotracker (blue) (Movie S1 with insets). We found that GFP-Rhes was enriched with globular mitochondria, which becomes positive for lysosome in a time dependent fashion (arrows, Movie S1). Intriguingly, enhanced GFP-Rhes signal intensity was associated with loss of mitotracker intensity, suggestive of mitochondrial degradation (arrow in GFP-Rhes). Collectively, these data indicated that Rhes associates with mitochondrial as well as lysosomal components in the intact striatum and in the cultured striatal neuronal cell line, where we found stronger association with globular mitochondria.

**Figure 1.**
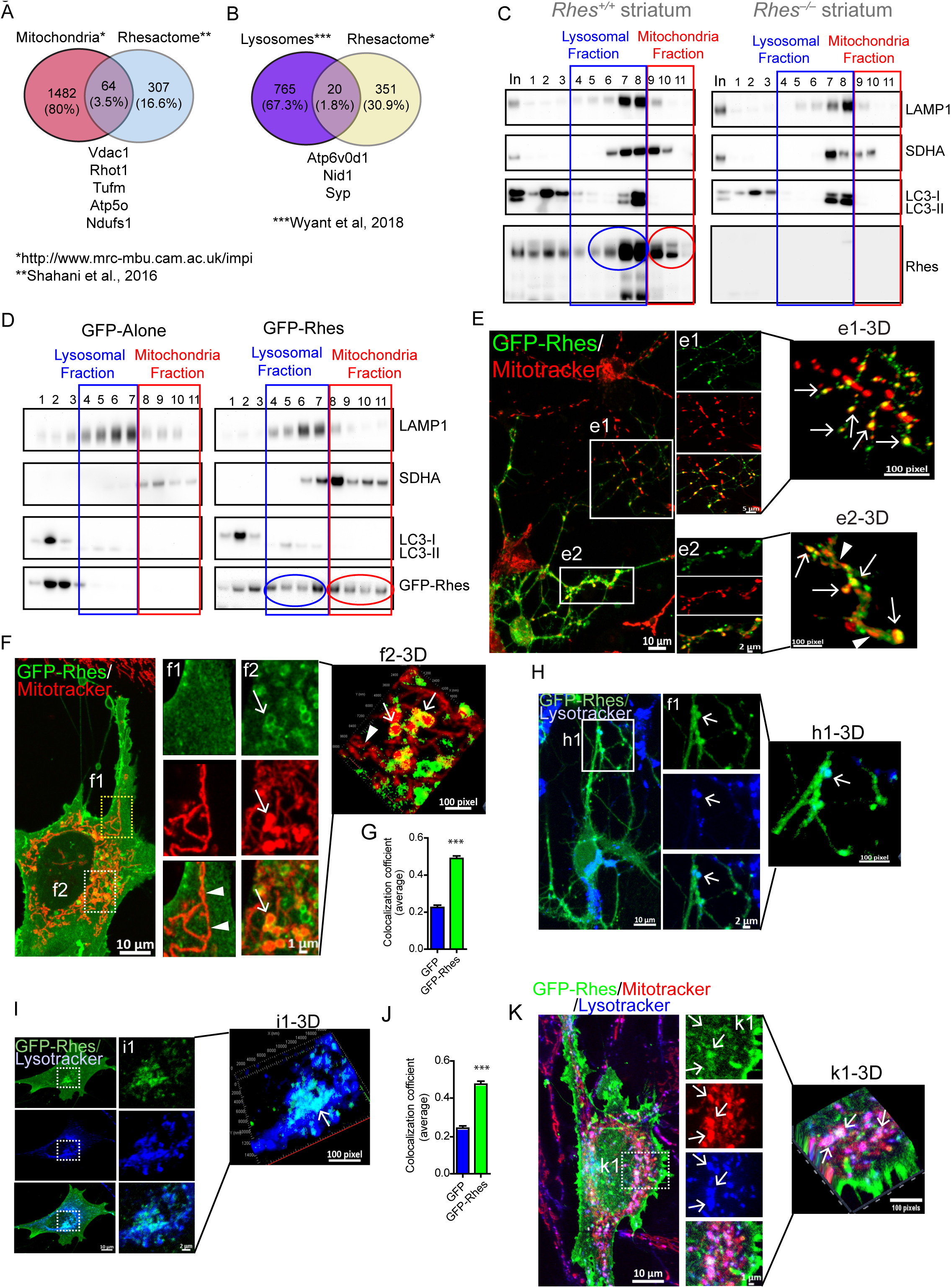
Rhes associates with mitochondria and lysosome in vitro and in vivo. (A, B) Venn diagram showing mitochondrial and lysosomal proteins found in Rhes interactome in vivo, “Rhesactome.” (C and D) Sucrose density gradient and Western blotting of striatal tissue (C) from Wt or Rhes KO mice, and striatal neuronal cells expressing GFP or GFP-Rhes (D) indicating lysosomal (blue) and mitochondrial (red) proteins across different fractions. Representative confocal images of primary striatal neuron (E), striatal neuronal cells (F) and their respective insets or 3D rendered images transfected with GFP-Rhes and co-stained with mitotracker orange (red). Arrow and arrowhead depict globular and elongated mitochondria respectively in neuronal processes. (G) Bar graph depicting average Pearson’s coefficient of colocalization (n=35-40 cells per group, Student’s-*t*-test, ***p<0.001. Data mean ± SEM. Representative confocal image of primary striatal neuron (H) or striatal neuronal cells (I) and their respective insets or 3D rendered image transfected with GFP-Rhes and co-stained with lysotracker (blue). Arrow represents the Rhes and lysosome colocalization. (J) Bar graph shows average Pearson’s coefficient of colocalization (n=39-42 cells per group, Student’s-*t*-test, ***p<0.001. Data mean ± SEM). (J) Confocal image, insets and 3D rendered image of striatal neuronal cells expressing GFP-Rhes co-stained with both mitotracker orange (red) and lysotracker (blue). Arrows indicates GFP-Rhes positive for mitochondria and lysosomes.

### Rhes affects basal mitophagy but not mitochondrial functions

Using cycloheximide (CHX) chase experiments, we tested whether Rhes affects basal mitophagy. As expected CHX treatment resulted in the degradation of SDHA, VDAC1, the mitochondrial markers, whose levels were further diminished by GFP-Rhes compared to GFP-alone overexpression (Fig. 2A). Note, levels of mTOR or Bip were not altered by CHX between the groups in these cells and downregulation of LC3-II was similar between GFP or GFP-Rhes expressing cells (Fig. 2B). Next, we wondered, whether Rhes can affect overall mitochondrial functions. To test this, we used seahorse assay, which measures the mitochondrial respiratory functions. We infected striatal neuronal cell lines with validated adenovirus empty (Ad-null) or adenovirus-Rhes (Ad-Rhes) (Swarnkar et al., 2015). As shown in Fig 2 C and D, Rhes expression did not affect the mitochondrial functions basally or in the absence or the presence of oligomycin (an inhibitor of ATP synthase), FCCP (uncouples oxidative phosphorylation), or rotenone (complex I inhibitor). Similarly, Rhes expression did not affect FCCP-induced mitophagy (Fig. 2, E and F). Thus, Rhes enhances basal mitophagy in CHX conditions, but may not affect the overall mitochondrial functions basally or in the presence of certain pharmacological modulators of mitochondria.

**Figure 2.**
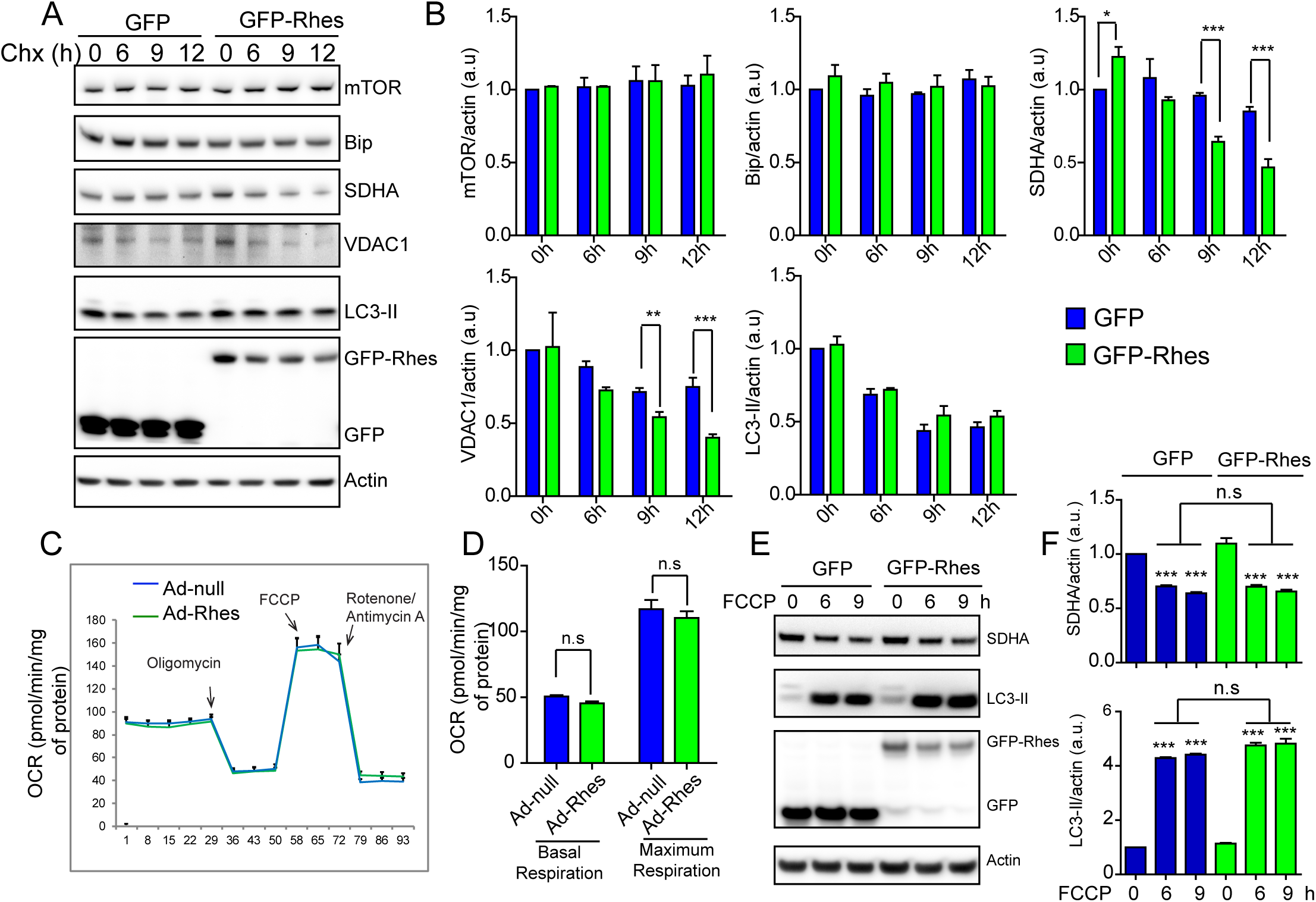
Rhes regulates basal mitophagy. (A) Representative western blots from GFP or GFP-Rhes transfected striatal neuronal cells treated with vehicle or cycloheximide (CHX, 100 µm) at indicated time point. Blots were probed with indicated antibodies. (B) Bar graph shows quantification of band intensities in panel A. Protein levels of mTOR or Bip or SDHA or VDAC1 or LC3-II in GFP or GFP-Rhes transfected cells were normalized to Actin. (n=3, Student’s-*t*-test, and; *p<0.05, **p<0.01, ***p<0.001. Data mean ± SEM). (C) Seahorse assay in striatal neuronal cells infected with Ad-null or Ad-Rhes for 48hr to assess basal oxygen consumption rate (OCR) prior to any injections and after injection of the complex IV inhibitor oligomycin, the uncoupler carbonyl cyanide 4-(trifluoromethoxy) phenylhydrazone (FCCP) and a combined injection of rotenone + antimycin A. (D) Bar graph shows the OCR at basal and maximum respiration for the indicated groups. (n = 6 per group). (E) Representative Western blot of indicated proteins and band intensity (F) from striatal neuronal cells expressing GFP or GFP-Rhes and treated with FCCP (1 µm) for the indicated time points. (n=3, One-way ANOVA test, ***p<0.001. n.s is not significant between indicated groups. Data mean ± SEM).

### Rhes diminishes mitochondrial functions and upregulates mitophagy in presence of 3-NP

To further assess the role of Rhes in mitochondrial functions, we investigated the effect of Rhes on 3-NP in seahorse assay. The rationale for testing 3-NP is twofold: a) 3-NP is known to promote lesion selectively in the striatum, and b) previously we found Rhes knockout mice are totally prevented from 3-NP induced striatal lesion compared to WT control (Mealer et al., 2013). In sea horse assay, as in Fig 2C, Rhes alone did not affect mitochondrial function in vehicle conditions. But, in presence of 3-NP, which diminished both the basal and maximal mitochondrial respiration, Rhes further potentiated the effect (Fig. 3A and B). This indicated that Rhes selectively worsens the 3-NP induced mitochondrial dysfunctions, but not oligomycin, FCCP or rotenone.

**Figure 3.**
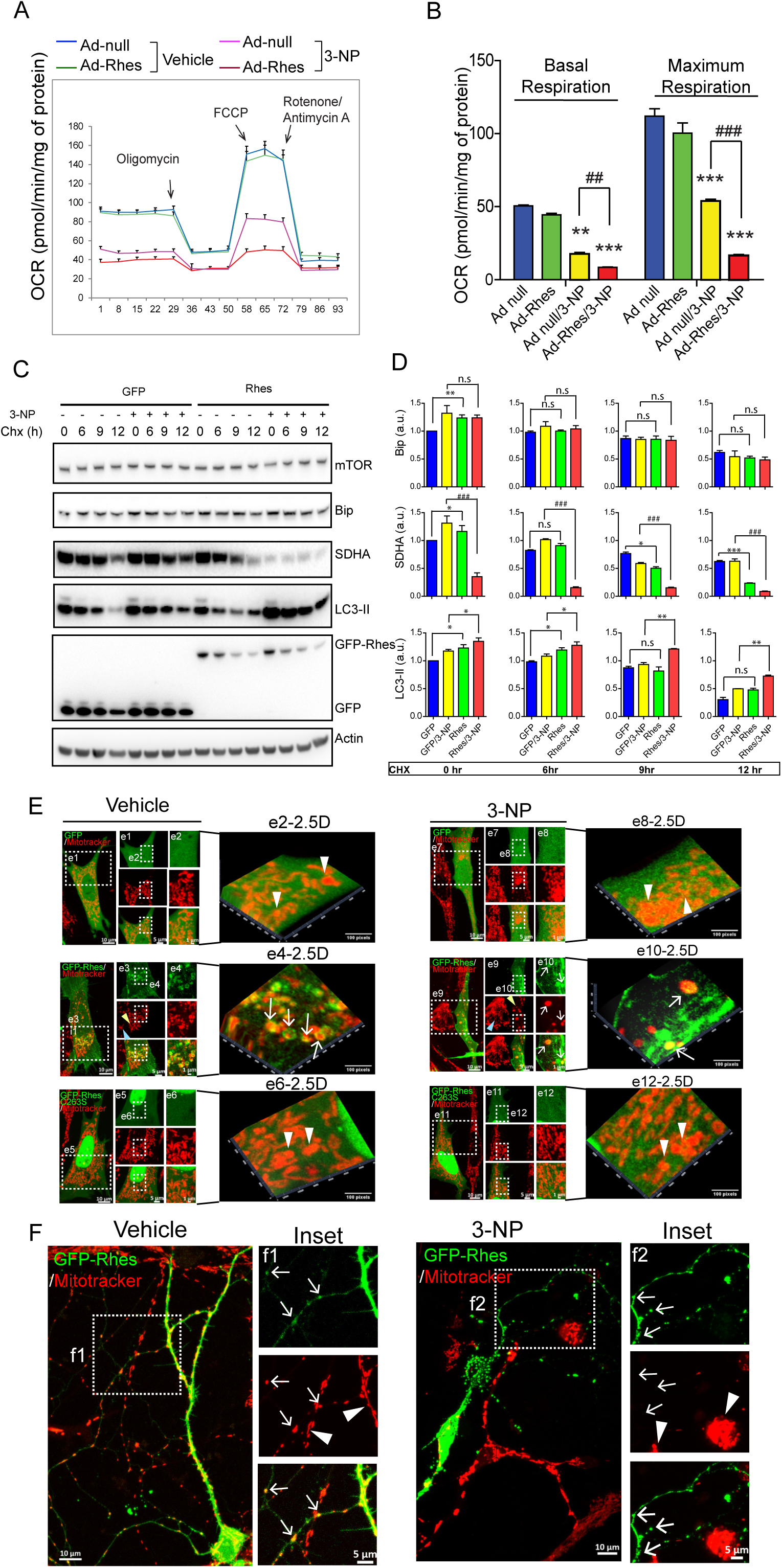
Rhes promotes mitophagy in presence of SDH inhibitor, 3-NP. (A) Shows seahorse assay in striatal neuronal cells infected with Ad-null or Ad-Rhes viral particles with vehicle or 3-NP (10 mM for 2 hr). As in Fig 2C, the graphs show the baseline OCR and OCR after oligomycin, FCCP and rotenone + antimycin A in vehicle or in 3-NP. (B) Bar graph shows the OCR measurements at basal and maximum respiration among indicated groups. [**p < 0.01; ***p < 0.001, compared to Ad-null and vehicle treated; ##p < 0.01; ###p < 0.001, between indicated groups, One-way ANOVA test (n = 6/group, data mean ± SEM)]. (C) Representative western blots of striatal neuronal cells transfected with GFP or GFP-Rhes, treated with vehicle or 3-NP (10 mM for 1hr). Samples were treated with CHX (100 µm) as indicated time points and blots were probed for indicated proteins. (D) Bar graph shows the quantification of indicated proteins [*p < 0.05, **p < 0.01, ***p < 0.001 vs GFP/vehicle and GFP-Rhes/Vehicle; ^###^p < 0.001 vs. GFP/3-NP and GFP-Rhes/3-NP (n=3, Data mean ± SEM; Student’s *t*-test). n.s. not significant. (E) Representative confocal images of striatal neuronal cells transfected with GFP or GFP-Rhes or GFP-Rhes C263S treated with vehicle or 3-NP and corresponding insets or 3D rendered images. Arrows indicate globular mitochondria positive for GFP-Rhes. Yellow and blue arrowhead indicate GFP-Rhes transfected cell and untransfected cell respectively in the same field. White arrowheads show globular mitochondria negative for GFP or GFP-Rhes C263S. (F) Representative confocal image of GFP-Rhes transfected primary striatal neuron and corresponding insets treated with vehicle or 3-NP. Cells were stained with mitotracker orange (Red). Arrow indicates GFP-Rhes positive neuronal processes colocalized with mitochondria in vehicle, but not in 3-NP treated neuron. Arrowheads indicate the mitotracker intensity is not affected in untransfected neighboring cell in the same field either in vehicle or 3-NP treatment.

To investigate the mechanisms, we assessed the effect of Rhes on mitophagy in striatal neuronal cell line or primary striatal neurons using biochemical and confocal imaging approaches. We used cycloheximide chase experiment to assess the mitophagy and Rhes markedly enhances mitophagy. As shown in Fig 4C, in GFP alone, there was a gradual loss of SDHA, both in vehicle and 3-NP conditions at 6, 9, and 12 hrs after CHX treatment. In presence of GFP-Rhes, loss of SDHA is more rapid in vehicle and almost all (>90%) of SDHA was eliminated in 3-NP, which is also accompanied by enhanced LC3-II production (Fig. 4 C and D). Next, we transiently expressed GFP, GFP-Rhes wt, GFP-Rhes C263S (mutant defective in membrane binding), in primary striatal neuronal cells by staining them with mitotracker and exposing them to vehicle or 3-NP (Fig. 4E). As expected, in vehicle treated neuronal cells, we found GFP-Rhes wt interacted with mitochondria (inset e4-2-5D, arrow) and that mitotracker staining was observed throughout the GFP-Rhes wt transfected (yellow arrowhead) and untransfected (blue arrowhead) neuronal cells (inset, e3). Contrary to this observation, in 3-NP treated neuronal cells, the dispersed mitotracker staining was completely disrupted in the GFP-Rhes transfected cells (inset e9, yellow arrowhead) but not in untransfected neuronal cells (inset e9, blue arrowhead). In GFP-Rhes transfected cells, the numerous globular mitochondria that were positive for Rhes were observed (e10-2.5D, arrow). Note, in GFP alone or GFP-Rhes C263S transfected cells the mitotracker signal appear undisrupted, and there no obvious association of GFP or GFP-Rhes C263S with mitochondria either in vehicle or 3-NP treated cells (e8-2.5D, e12-2.5D, arrowhead). We further analyzed the mitotracker intensity by flowcytometry in striatal neuronal cells, infected with either Ad-null or Ad-Rhes, treated with vehicle or 3-NP. We found that mitotracker intensity was reduced by 30% in Ad-Rhes infected cells compared to Ad-null infection after 3-NP treatment (Fig. S2), indicating Rhes robustly diminishes mitochondrial intensity upon 3-NP. Analogous to striatal cell lines in primary striatal neurons also we found GFP-Rhes wt interacted with mitochondria (Fig. 3F, inset f1, arrow) and that mitotracker staining was observed throughout the GFP-Rhes wt transfected and untransfected primary neuronal processes (inset, f1 arrowhead). However, in 3-NP treated neuronal cells, the mitotracker staining was almost completely abolished in the GFP-Rhes wt transfected cells (inset f2, arrow) but not in untransfected neuronal cells (inset f2, arrowhead). Note, GFP alone (control) transfected primary neuron does not show reduction in mitotracker staining after 3-NP treatment (Fig. S3). Together, this data indicates Rhes exacerbates mitochondrial dysfunction and promotes mitophagy in presence of SDHA inhibitor, 3-NP.

**Figure 4.**
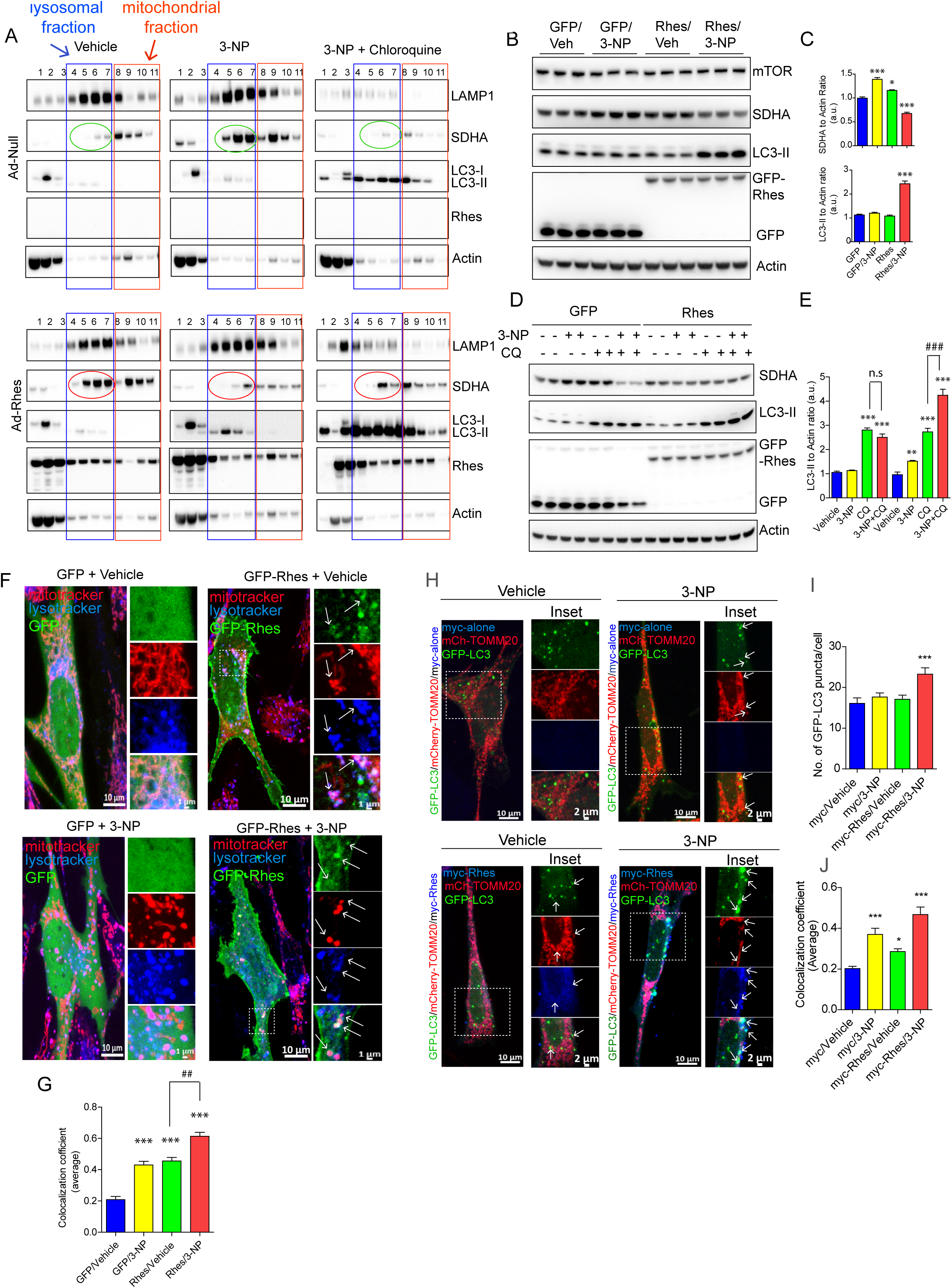
Rhes increases the mitophagy flux. (A-E) Representative immunoblots sucrose gradient (A) infected with Ad-null or Ad-Rhes or whole cell lysate (B and D) from GFP or GFP-Rhes transfected striatal neuronal cells treated with vehicle or 3-Nitropropionic acid (3-NP) for 2 h, with or without cholorquine (50 µM), probed for indicated proteins. (C and E) Bar graph shows the quantification of SDHA and LC3-II protein level normalized to actin. [(*p < 0.05; ***p < 0.001 compared to GFP/Veh (Vehicle) (n=6, Data mean ± SEM; One-way ANOVA test)]. (F) Representative confocal image of striatal neuronal cells transfected with GFP or GFP-Rhes, treated with vehicle or 3-NP (10 mM for 2 hr). Cells were stained with mitotracker (Red) and lysotracker (Blue). Inset shows the magnified region from selected area and arrows represents the globular mitochondria positive for GFP-Rhes and lysotracker. (G) Bar graph shows the average of Pearson’s coefficient of colocalization between mitotracker and lysotracker in GFP or GFP-Rhes transfected cells (n=21-28 cells per group, One-way ANOVA vs. GFP/vehicle; ***p<0.001. ^##^p<0.01 between Rhes/vehicle vs. Rhes/3-NP using Student’s *t*-test. Data mean ± SEM. (H). Representative confocal images of striatal neuronal cells co-transfected with GFP-LC3 and mCherry-TOMM20 along with either myc-alone or myc-Rhes. Cells were treated with Vehicle or 3-NP (10 mM for 2 hr). Cells were fixed and stained for myc antibody (blue). Corresponding inset shows the magnified area from selected region. Arrows show the colocalization between GFP-LC3 and mCherry-TOMM20 in indicated groups. (I-J) Bar graph shows (I) the quantification of GFP-LC3 puncta per cell and (J) average of Pearson’s coefficient of colocalization between GFP-LC3 and mCherry-TOMM20 in indicated groups. n=21-24 cells per group; One-way ANOVA, ***p<0.001. Data mean ± SEM.

### Rhes promotes mitophagy flux in presence of 3-NP

Next, we biochemically estimated whether Rhes alters the mitophagy flux in striatal neuronal cell lines in gradient fractionation (Fig. 4A) and in total lysate (Fig. 4, B-E). In Ad-null condition (control), as expected we found an accumulation of SDHA in the mitochondrial fraction in vehicle treated groups (Fig. 4A). Addition of 3-NP resulted in a robust accumulation of SDHA in the lysosomal fractions (compare green circles). Blocking autophagy with chloroquine (CQ), however, resulted in a diminished accumulation of SDHA and enhanced LC3-II levels in control (Fig. 4A), indicating that 3-NP might induce new mitochondrial production, in control condition. In Ad-Rhes, however, higher levels of SDHA were found in vehicle treated conditions. Treatment of 3-NP in Ad-Rhes condition resulted in a reduction of SDHA levels in the lysosomal fractions (compare red circles). Co-treatment with CQ showed an increased SDHA levels in lysosomal fraction, which is also accompanied by a rapid upregulation of LC3 conversion (Fig. 4A). Together biochemical fraction data suggest that Rhes upregulates mitophagy flux in 3-NP condition.

In total lysate, both in vehicle treated GFP or GFP-Rhes expressing cells, we found there was no difference in SDHA levels (Fig. 4, B and C). Upon 3-NP, however, there was an increased SDHA levels in GFP expressing cells, but in GFP-Rhes expressing cells the SDHA levels were diminished, which is also accompanied by a rapid upregulation of autophagy, as measured by increased LC3 conversion (Fig. 4, B and C). Addition of CQ has further increased the SDHA and LC3-II levels in 3-NP condition, as in 4A (Fig. 4, D and E). Together these data indicate that Rhes increases autophagosomes and lysosomes accumulation around mitochondria and mediates mitophagy flux in the presence of 3-NP. Similarly, in immunocytochemistry, SDHA staining is also diminished in GFP-Rhes transfected striatal neuronal cells, treated with 3-NP compared to vehicle treatment (Fig. S4). Next, we investigated whether Rhes recruit lysosomes to globular mitochondria using lysotracker and GFP-LC3 by confocal microscopy. As predicted, we readily found GFP-Rhes localization with mitotracker and lysotracker in presence of 3-NP, and there was enhanced localization of GFP-Rhes with lysotracker in presence of 3-NP (arrow, Fig. 4, F and G). Similarly, there was an enhanced number of GFP-Rhes/LC-3 puncta (arrow, Fig 4, H and I) and their co-localization with mCherry TOMM20, a mitochondrial marker, by 3-NP (Fig. 4J).

### Rhes promotes the formation of mitophagosomes *in vivo*

Since Rhes interacted with globular mitochondria and lysosomes (Fig. 1), and also modulates basal mitophagy which is further upregulated in 3-NP (Figs. 2-4), we posited that Rhes may activate mitophagy *in vivo*. To test this thoroughly in vivo, we carried out behavioral, pathological and ultrastructural analysis (using transmission electron microscopy) of the striatum in WT and Rhes KO mice after systemic injection of 3-NP (see experimental scheme Fig 5A). Consistent with our earlier report (Mealer et al., 2013), 3-NP administration to WT mice promoted motor abnormalities that was diminished in Rhes KO mice (Fig. 5B-E). 3-NP also induced striatal specific lesion in WT striatum, which was completely prevented in the Rhes KO striatum (Fig. 5F, arrowhead) (Mealer et al., 2013). Microscopic analysis of the 3-NP induced striatal-specific lesions found pyknotic nuclei and a diminished neuronal staining in WT mice, while KO striatum displays healthy nuclei and normal neural staining (Fig. 5F, compare WT & KO, insets f2)(Mealer et al., 2013). Thus, 3-NP promotes lesions selectively in the striatum of the WT mice but not in the Rhes KO striatum, a highly reproducible phenotype (Mealer et al., 2013). To address why Rhes KO are resistant to 3-NP induced lesion, we hypothesized that ultrastructural analysis might provide a clue. We carried out time-lapse ultrastructural analysis using serial cortico-striatal section electron microscopy of 2^nd^ and 3^rd^ day after systemic administration of 3-NP. On the 2^nd^ day, we found no gross changes in the overall cell morphology or mitochondria ultrastructure between WT and KO, which were similar to control (Fig. 5G and insets). On the 3^rd^ day, in WT, we found neuronal shrinkage and abnormal, swollen mitochondria with broken inner membrane and matrix in the striatum (Fig. 5G, arrowhead). And, those abnormal striatal neuronal changes were completely absent in the Rhes KO (Fig. 5G). Mitochondrial circularity index analysis (length/breadth) showed that the frequency of circular mitochondria is high in Rhes WT compared to Rhes KO neuron after 3-NP treatment (Fig. 5I). Cortical neurons from either genotype showed no striatal lesion (arrowhead) or mitochondrial abnormalities (insets, a1 and b1), indicating that 3-NP induced striatal lesion (arrow) and mitochondrial abnormalities (inset b2) is selective to the striatum and it requires Rhes (Fig. S5, A and B). Further examination of ultrastructure revealed numerous mitophagosomes, where the mitochondria with cristae debris were engulfed within the double layer membranes resembling autophagosomes (Fig. 5J, and arty rendering) and there was a diminished frequency of number of healthy-looking mitochondria, but enhanced swollen mitochondria and mitophagosomes in Rhes WT striatum, compared Rhes KO (Fig. 5K). Thus, Rhes KO prevents the formation of neuronal shrinkage, mitochondrial swelling, and induction of mitophagosomes by 3-NP. All together, these data indicated that Rhes physiologically regulates 3-NP induced mitochondrial dysfunction and promotes the formation of globular mitochondria and mitophagosomes, which coincides with worsening motor phenotype *in vivo*.

**Figure 5.**
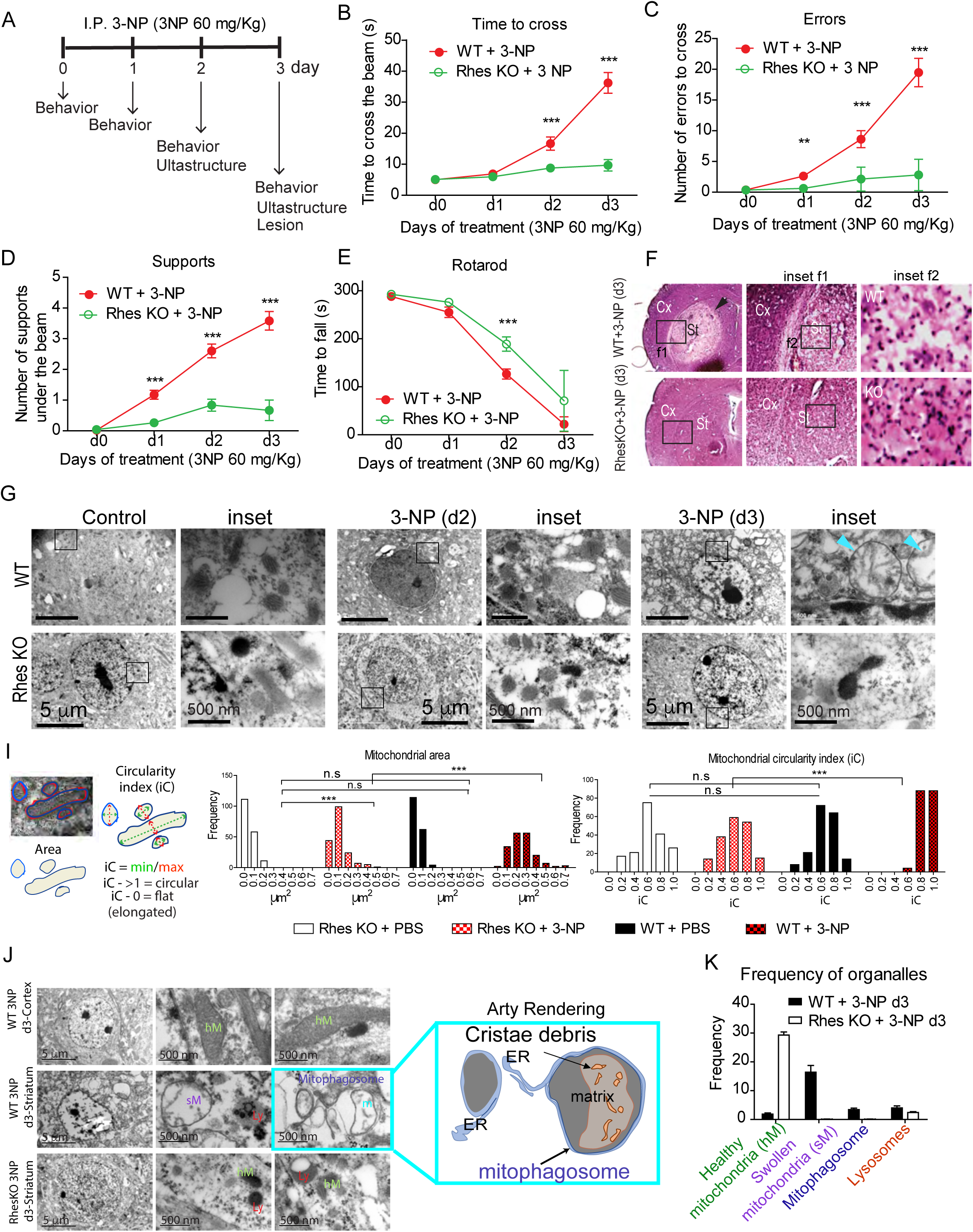
Rhes promotes mitophagy in vivo. (A) Experimental design for 3NP treatment in mice. (B-D) Beam walk analysis during 3NP treatment for WT (red circles) and Rhes-KO (green circles) groups. (E) Changes in the latency to fall for the rotarod test during 3NP treatment. Two-way Anova post hoc test. (F) Representative hematoxylin-eosin (H&E) stained sections for WT and Rhes-KO mice after 3 days of 3NP treatment. Representative electronic micrographs at low (G) and high (H) magnification at day 2 (d2), day 3 (d3) during 3NP treatment, insets in G are shown in high magnification, H. (I) Ultrastructural changes analysis using frequency histograms of the mitochondrial area and circularity index (iC) for the WT and Rhes KO groups treated with vehicle (PBS) or 3NP. One-way ANOVA analysis of variance (Bartlett’s test), Bonferroni’s posthoc. *** p < 0.001. (J) Representative electronic micrographs from striatal neurons, mitochondria and lysosomes after 3 days of 3NP treatment. (K, upper panel) Schematic representation of mitophagosome components based on the indicated micrograph in J. (K, bottom panel), Quantification of the healthy and swollen mitochondria, lysosomes and mitophagosomes frequency at third day of 3NP treatment in WT and Rhes KO mice. Graphics and statistics were made using PRISMA-GraphPad software, *** p < 0.001, ** p < 0.01, determined by two-way ANOVA, Bonferroni’s *post hoc*.

### Rhes diminishes mitochondrial potential (ΔΨ*m*) and promotes cell death in the presence of 3-NP

To understand the mechanisms, and because Rhes can interact with VDAC1 (Fig. 1A), which is a component of membrane permeability pore, we rationalized that Rhes may affect mitochondrial potential (ΔΨ*_m_*_)_. To investigate this, we tested whether Rhes alters ΔΨ*_m_* using tetramethylrhodamine (TMRM), a widely used cell permeable fluorescent dye that binds mitochondria with intact ΔΨ*_m_.* We infected striatal neuronal cell lines with Ad-null or Ad-Rhes (Swarnkar et al., 2015) for ∼32 hours and added TMRM. We quantified TMRM signal intensity using FAC sorting in vehicle or 3-NP at 15 min, 30 min and 2hr (Fig. 6A). Notably, TMRM (red) signals were not altered in vehicle condition both in Ad-null or Ad-Rhes expressing cells. But in the presence of 3-NP the TMRM signal intensity is rapidly diminished only in Ad-Rhes expressing cells but not Ad-null in a time-dependent manner, indicating a rapid loss of ΔΨ*_m_* by Rhes (Fig. 6A). This effect is specific to Rhes wt, as GFP-Rhes C263S mutant failed to alter TMRM signal, which is like GFP control (Fig. 6B, C). Next, as Rhes is required for 3-NP induced striatal lesion in vivo [(Mealer et al., 2013), and Fig 5], we investigated whether Rhes modulates striatal cell death in vitro, using propidium iodide (PI) staining, a fluorescent dye that enters cells only when they are compromised (Subramaniam et al., 2004), and FAC sorting. As shown in Fig. 6D, we found Ad-null control expressing cells did not show changes in PI+ cells either in vehicle or 3-NP condition (∼5-7%). Similarly, Ad-Rhes expressing cells in vehicle did not show changes in PI+ cells (∼5%). However, in Ad-Rhes and 3-NP condition, there was robust increase in PI+ cells (∼25%, Fig. 6E). To further establish that cell death effect is specific to Rhes wt, we FAC sorted GFP alone, GFP-Rhes wt and GFP-Rhes C263S cells and added vehicle or 3-NP (See scheme. Fig 6F). As shown in the Fig 6, G and H, Rhes wt showed ∼27% cell death as measured by PI staining in presence of 3-NP, whereas GFP or GFP-Rhes C263S, showed ∼ 5-10% cell death (Fig. 6H). Collectively these data indicate Rhes diminishes ΔΨ*_m_* and promotes cell death in the presence of 3-NP.

**Figure 6.**
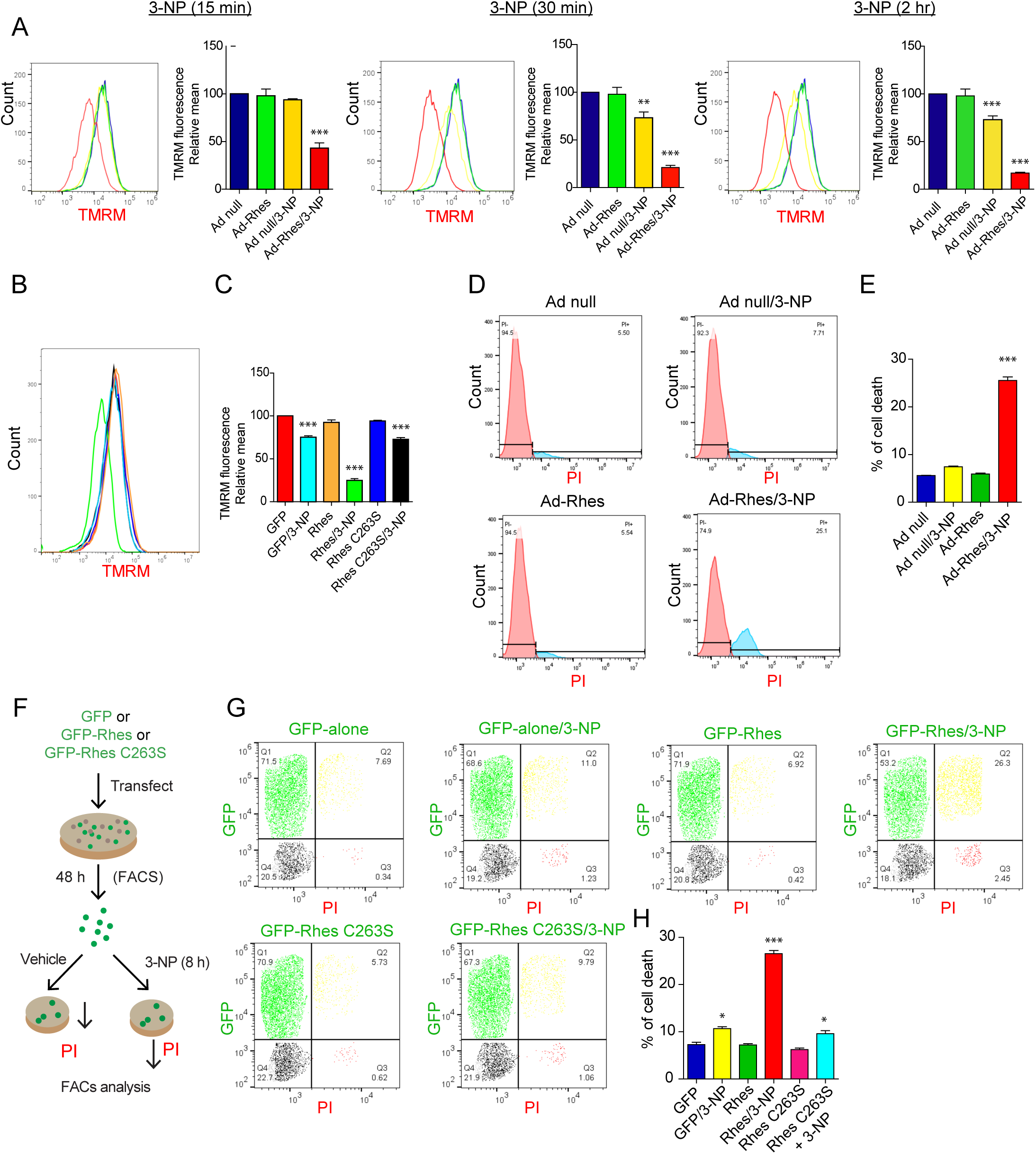
Rhes diminishes mitochondrial membrane potential (ΔΨ*_m_*) and promotes cell death. (A) Representative flowcytometry plot of striatal neuronal cells infected with Ad-null or Ad-Rhes in presence of vehicle or 3-NP (10 mM, for 15 min, 30 min or 2 hr). Cells were stained for TMRM (ΔΨ***_m_*** indicator) and analyzed in flowcytometry. Bar graph shows the relative TMRM mean intensity. **p < 0.01, ***p < 0.001 vs. Ad-null and vehicle (n = 3/group; Data mean ± SEM; one-way ANOVA). (B) Flowcytometry plot of striatal neuronal cells transfected with GFP or GFP-Rhes or GFP-Rhes C263S treated with vehicle or 3-NP (10 mM and for 2 hr) and stained for TMRM. (C) Bar graph shows the relative TMRM mean intensity. ***p < 0.001 vs. GFP/Vehicle sample (n = 3 per group; Data mean ± SEM; One-way ANOVA). (D) Flowcytometry plot of striatal neuronal cells infected with Ad-null or Ad-Rhes treated with vehicle or 3-NP (10 mM for 8 hr), stained with propidium iodide (PI). (E) Bar graph shows the % of cell death in indicated groups. ***p < 0.001 vs. Ad-null/Vehicle (n = 3 per group; Data mean ± SEM; One-way ANOVA) (F) *Experimental design* for panel G. (G) Flowcytometry plot of FACs sorted striatal neuronal cells positive for GFP or GFP-Rhes or GFP-Rhes C263S treated with vehicle or 3-NP (10 mM and for 8 hr) and stained for propidium Iodide (PI). (H) Bar graph shows the % of cell death in indicated group. *p < 0.05, ***p < 0.001 vs. GFP/Vehicle sample (n = 3 per group; Data mean ± SEM; One-way ANOVA).

## Rhes promotes mitophagy, disrupts ΔΨ*_m_* and promotes cell death via Nix

To investigate the mechanisms by which Rhes may activate mitophagy, we considered whether Rhes interacts with well-known mitophagy modulators, such as Parkin, PINK1, DRP1 or Nix (Jin and Youle, 2012). In an affinity purification experiment, GST-Rhes did not interact with Parkin but it readily interacted with mTOR (Fig. 7A), as shown in our previous report (Subramaniam et al., 2012). We did not observe interaction of Rhes with either DRP1 or Pink1 (Fig. S6, A and B). However, GST-Rhes readily interacted with Nix, a known mitophagy receptor (Novak et al., 2010) (Fig. 7B, C). Rhes interaction with overexpressed Nix (Fig. 7B) or endogenous Nix (Fig. 7C), increase with 3-NP treatment, however Rhes interaction with mTOR remain unaltered (Fig. 7C). GST-Rhes is also affinity purified with LC3 without or with 3-NP and showed a consistent and enhanced interaction with Nix in presence of 3-NP (Fig. 7D). Next, we found that the C-terminal SUMO E3 ligase domain (171-266 aa), but not the N-terminal GTPase (1-181 aa), or the fragment (147-216 aa) interacted strongly with Nix (Fig. 7E). Additionally, we found membrane binding defective mutant Rhes C263S, but not GTP-binding mutant, fails to interact with Nix (Fig. S6C). Interaction of Rhes with mTOR, on the other hand, seems to be more towards the N-terminal side of Rhes (Fig. 7E). Thus, Rhes associates strongly with Nix in the presence of 3-NP and the binding requires intact membrane binding domain.

**Figure 7.**
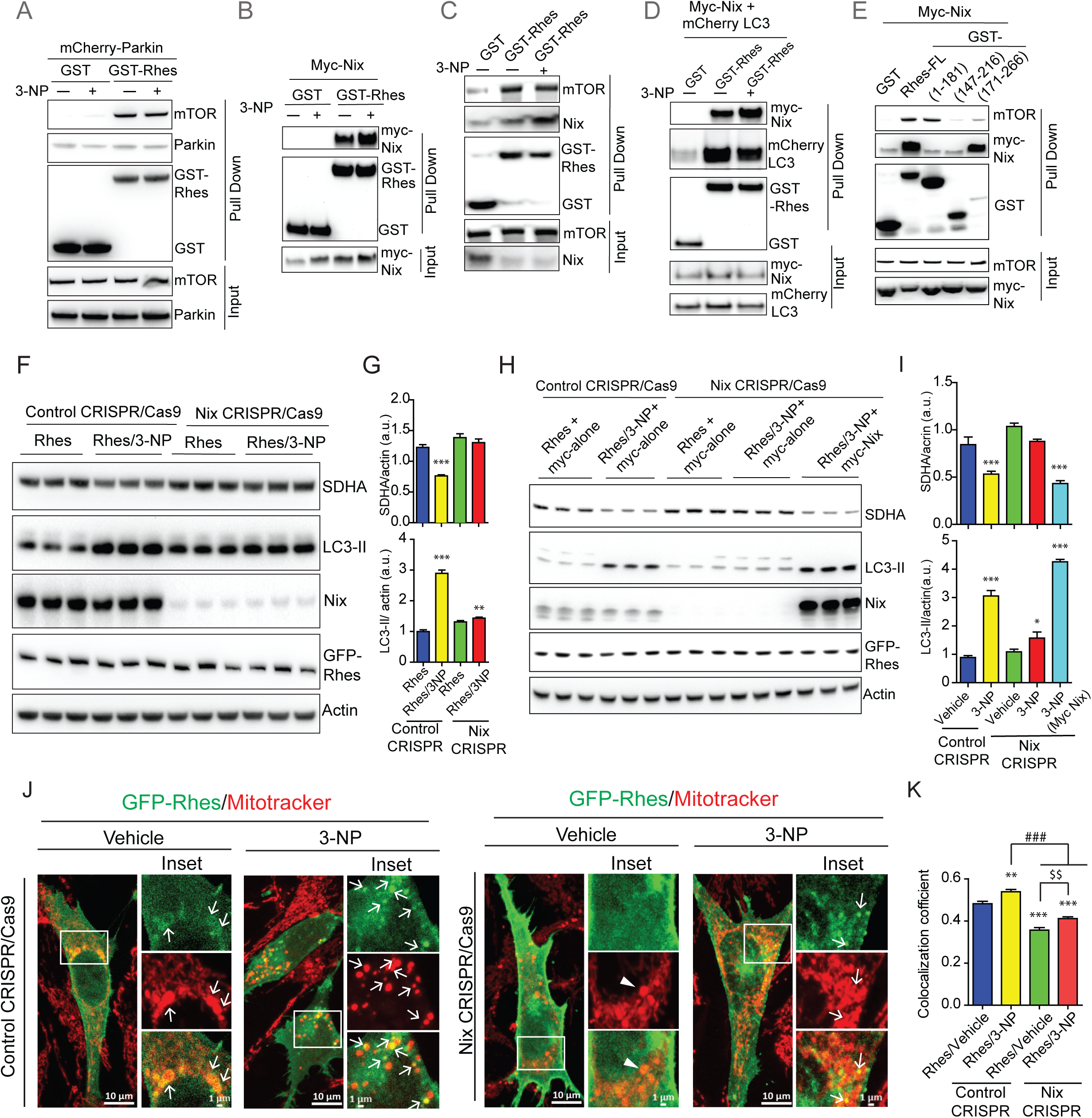
Rhes interacts with Nix and promotes mitophagy. (A-E) Representative Western blot of indicated proteins after glutathione-affinity purification of HEK 293 cells transfected with GST or GST-Rhes (Full length-FL or fragments), mCherry-Parkin, myc-Nix or mCherry-LC3 plasmids that are exposed to vehicle or 3-NP (10 mM, 2 hr) wherever indicated. Input 5% of total lysate. (F) Representative Western blot of control or CRISPR/Cas9-mediated Nix depleted striatal neuronal cells that were transfected with GFP or GFP-Rhes and treated with vehicle or 3-NP (2 hr). (G) Bar graph shows the quantification for SDHA and LC3-II. **p < 0.01, ***p < 0.001 vs. control CRISPR/vehicle (n=4, Data mean ± SEM; One-way ANOVA). (H) Representative Western blots of indicated proteins from control CRISPR or NIX CRISPR/Cas9 striatal cells transfected with GFP-Rhes and myc-alone or myc-Nix constructs that are treated with vehicle or 3-NP (10 mM for 2 hr). (I) Bar graph shows normalized SDHA and LC3-II protein levels. *p < 0.05; ***p < 0.001 vs. control CRISPR/vehicle (n=4, Data mean ± SEM; One-way ANOVA). (J) Representative confocal images and their insets of control CRISPR/Cas9 or Nix CRISPR/Cas9 striatal neuronal cells transfected with GFP-Rhes and co-stained for mitotracker (red) that were treated with vehicle or 3-NP (10 mM for 2 hr). Arrows represents the globular mitochondria, positive for GFP-Rhes. (K) Bar graph shows the average of Pearson’s coefficient of colocalization between GFP-Rhes and mitochondria in indicated groups. [(n=33-37 cells per group; ***p<0.001, One-way ANOVA; ^###^p<0.001, ^$$^p<0.01, Student’s *t*-test. Data mean ± SEM].

Because Rhes interacted strongly with Nix, we hypothesized that Rhes may promote mitophagy via Nix. To test this hypothesis, we depleted endogenous Nix in the striatal cells using CRISRP/Cas-9 tools. While GFP-Rhes promoted a significant loss of SDHA and LC3 productions in CRISPR/Cas-9 control cells, it completely failed to do so in CRISRP/Cas-9 Nix-depleted cells (Fig. 7, F and G), which resulted in ∼90% depletion of Nix (Fig. S7A). This finding indicated Nix is essential for Rhes-induced mitophagy. Furthermore, when we reconstituted Nix in Nix depleted cells Rhes was able to restore mitophagy and increase LC-3 conversion (Fig. 7, H and I), suggesting that Nix is critical for Rhes-mediated mitophagy. Consistent with this, in confocal data, we found that while in wt control cells, Rhes promoted rapid loss of mitotracker intensity that is accompanied by formation of numerous globular mitochondria (arrow) (Fig. 7J). In Nix depleted cells, we found a) Rhes was unable to interact with globular mitochondria (Fig. 7J, arrowhead), b) upon 3-NP, a very few Rhes colocalization with mitochondria were seen and c) most of the mitotracker signal intensity remains unaltered by 3-NP. Pearson’s colocalization coefficient further supported that Rhes and mitochondria interacts less in Nix depleted cells compared intact cells (Fig. 7K). Together this data suggests that Rhes interacts with globular mitochondria and the promotes mitophagy via Nix.

Next, as Rhes decreases ΔΨ*_m_* in presence of 3-NP, we wondered whether this process requires Nix. As shown in Fig 8 A and B, while Rhes diminishes TMRM signals in control cells and this effect is not observed in Nix depleted cells. Similarly, the observed Rhes-induced cell death in presence of 3-NP in control cells was also markedly diminished in Nix depleted cells (Fig. 8C and D). Moreover, reconstitution of Nix significantly elevated Rhes-mediated cell death in 3-NP (Fig. 8D). Nix is localized to endoplasmic reticulum (ER) as well as mitochondria (Mughal et al., 2018). We used ER-targeted, Nix where transmembrane (TM) domain of Nix was replaced by CytoB and mitochondria-targeted Nix, where TM is replaced with monoamine oxidase located in the outer mitochondrial membrane (Mughal et al., 2018). We found Rhes affinity with both ER and mitochondrial targeted Nix (Fig. 8, E and F). However, in a reconstitution FACS/TMRM experiment, we found replenishing only the mitochondria-targeted Nix, but not ER-targeted Nix, restores the loss of TMRM signal by Rhes (Fig. 8, G and H). Collectively, this data indicated that Rhes interacts with both ER-and mitochondrial-targeted Nix and disrupts ΔΨ*_m_* mostly via the later.

**Figure 8.**
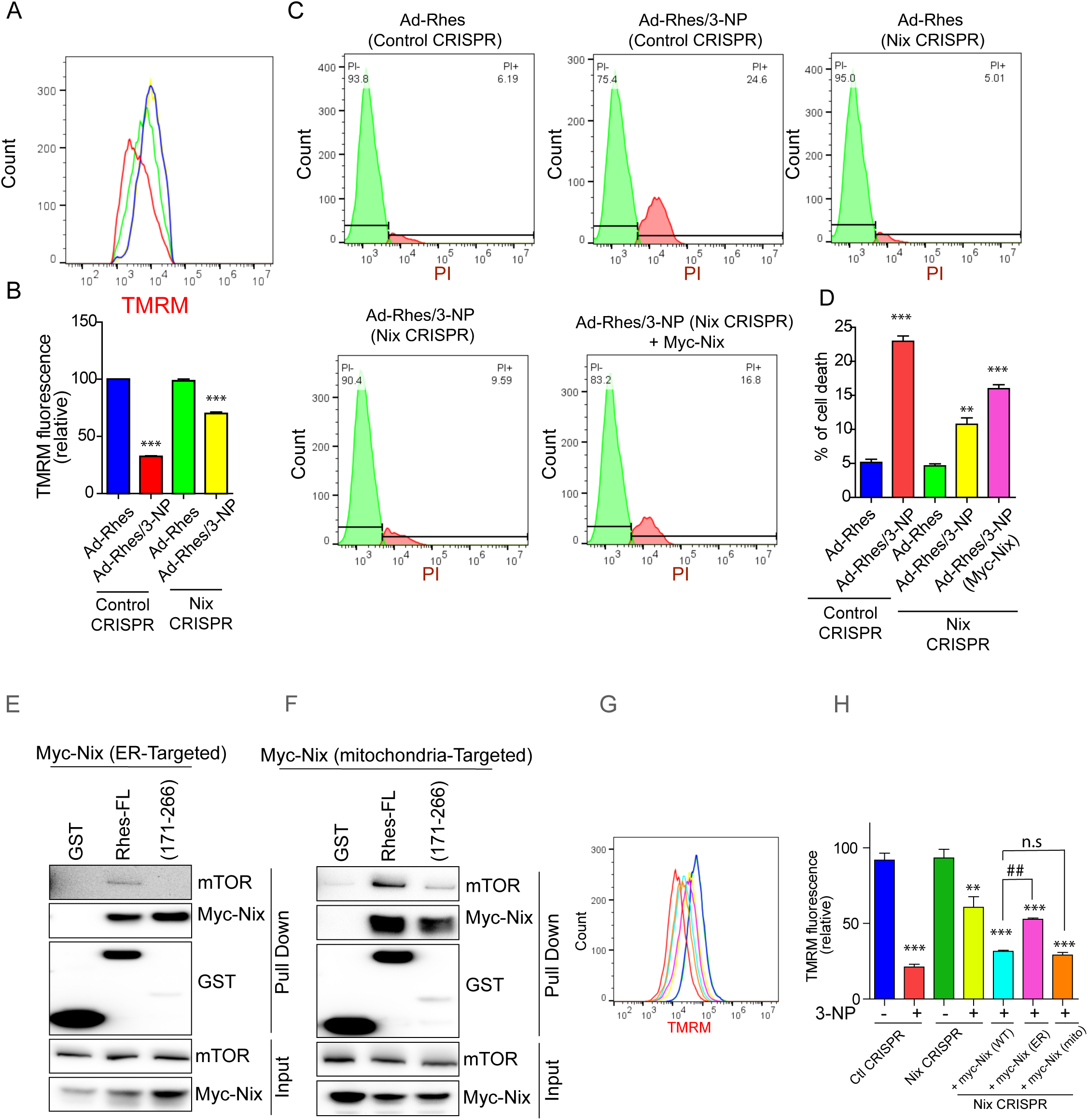
Rhes promotes cell death and disrupts mitochondrial potential (ΔΨ*_m_*) via mitochondrial targeted Nix. (A) Flowcytometry plots of control CRISPR/Cas9 and or Nix CRISPR/Cas9 striatal neuronal cells, either infected with Ad-null or Ad-Rhes viral particles. (B) Bar graph shows the relative TMRM fluorescence intensity. ***p < 0.001(n = 3 per group; Data mean ± SEM; One-way ANOVA). (C) Flowcytometry plot of control CRISPR/Cas9 and or Nix CRISPR/Cas9 striatal neuronal cells infected with Ad-null or Ad-Rhes treated with vehicle or 3-NP (10 mM for 8 hr) that are stained with PI. Nix CRISPR/Cas9 striatal neuronal cells reconstituted with Nix by transfecting myc-Nix cDNA. (D) Bar graph shows the % of cell death in indicated groups. ***p < 0.001, n = 3 per group; Data mean ± SEM; One-way ANOVA). (E-F) Representative GST pull down and western blots in HEK293 cells transfected with GST or GST-Rhes or GST-Rhes 171-266 plasmid together with either myc-Nix-ER (targeted to ER) (E) or myc-Nix-mitochondria (targeted to mitochondria) (F). Input (5%) of total lysate. (G) Flowcytometry plot and (H) Bar graph of striatal neuronal cells infected with Ad-Rhes and transfected with indicated vectors showing TMRM fluorescence signal in vehicle or 3-NP (10 mM for 2 hr) treated striatal neuronal cells (control or Nix depleted cells). n = 3/group, **p < 0.01, ***p < 0.001, ^##^p < 0.01, Data mean ± SEM; One-way ANOVA. Not significant (n.s).

### Rhes travels from cell-to-cell and interacts with the globular mitochondria in the neighboring cells via Nix

We recently reported Rhes induces the biogenesis of tunneling (TNT)-like cellular protrusion, “Rhes tunnel,” through which Rhes travels from cell-to-cell (Sharma and Subramaniam, 2019). It is unknown whether intercellular signaling can regulate mitochondrial turnover. So, we considered whether Rhes can interact with damaged mitochondria in the neighboring cells. We designed an experiment in which we co-cultured FACS sorted GFP-Rhes expressing cells (donor cells) with the acceptor cells that were exposed to 3-NP or vehicle (acceptor cells), followed by confocal imaging with mitotracker (see scheme Fig 9A). As expected, we found GFP-Rhes positive TNT-like protrusions (closed arrow) and numerous GFP-Rhes (open arrow) in the neighboring cell in both vehicle and 3-NP treated cells (Fig 9B). Interestingly, we found numerous GFP-Rhes puncta that were colocalized with globular mitochondria in cell that are exposed to 3-NP (arrowhead) in control acceptor cell. But when GFP-Rhes donor cell are cocultured with Nix depleted acceptor cells, we do observe TNT-like protrusions (closed arrow) and numerous puncta (open arrow), but we failed to observe GFP-puncta localization with globular mitochondria in the Nix-depleted acceptor cell (Fig 9B). Pearson’s colocalization coefficient further confirmed a marked reduction in Rhes localization with mitochondria in Nix depleted cells (Fig. 9C). This indicates that Rhes can travel from cell-to-cell and interact with damaged mitochondria via Nix. Thus, this data raises a possibility that Rhes can act as an intercellular mitochondrial surveillance factor.

**Figure. 9.**
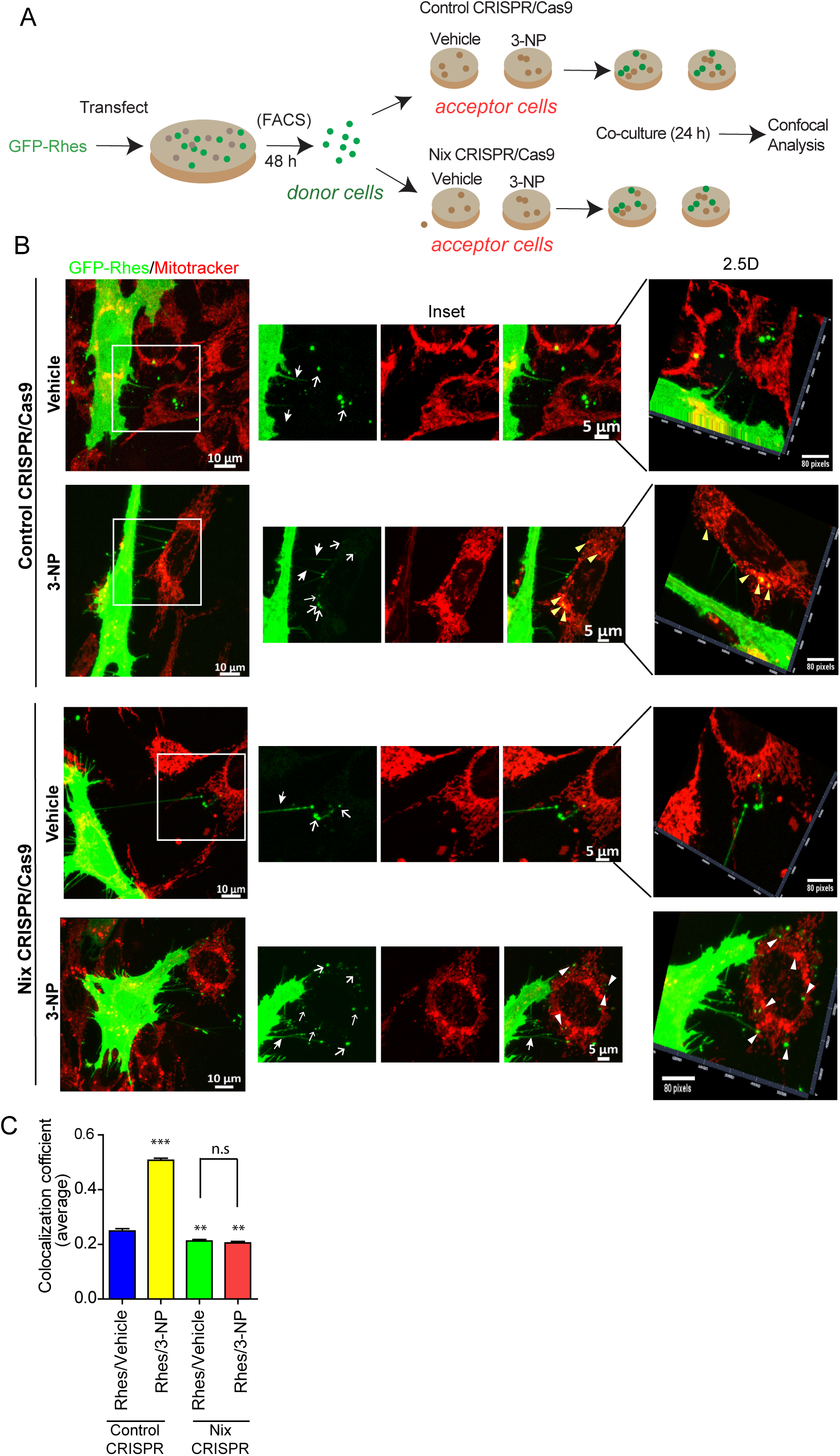
Rhes travels Intercellularly and interacts with damaged mitochondria via Nix. (A) Experimental design for panel B. (B) Representative confocal image of GFP-Rhes (FAC sorted) striatal neuronal cells (*donor cells*) cocultured with vehicle or 3-NP treated control or Nix depleted (Nix CRISPR) cells (*acceptor cells*). Insets show the magnified region from selected area. Corresponding insets were proceeded for 2.5-dimensional rendering. Closed arrow indicates Rhes-induced TNT-like protrusions. Open arrow represents GFP-Rhes puncta in *acceptor cell*. Yellow and white arrowhead indicates presence or lack of colocalization of GFP-Rhes with mitotracker, respectively. (C) Bar graph shows average of Pearson’s coefficient of colocalization between GFP puncta and mitotracker in neighboring *acceptor cells* where Rhes is transported from *donor cell*. n=72-90 GFP-Rhes puncta/group were counted from 13-18 cells per group. **p<0.01, ***p<0.001, One-way ANOVA. Not significant (n.s).

## Discussion

To date, the brain tissue specific regulator(s) of mitophagy or its role in neuronal vulnerability remains less understood. By using neuronal and mouse model combined with electron and live-cell confocal microscopy, we have systematically investigated role of Rhes in mitophagy. For the first time, to our knowledge, we found that striatal enriched protein in the promotion of mitophagy and that it requires Nix, a known mitophagy receptor (Novak et al., 2010). Despite abundant and comparable expression of Nix both in the striatum of WT and Rhes KO (Fig, S7B) the removal of damaged mitochondria via mitophagy requires Rhes demonstrating its “mitophagy ligand-like” capabilities in vivo. In addition, this study demonstrate the role of Rhes in intercellular surveillance of mitochondria through the cellular protrusion that we recently discovered (Sharma and Subramaniam, 2019) and lay a foundation for understanding the unprecedented complexity by which Rhes may signal mitophagy within and outside the striatum.

Half-lives of mitochondria in rat whole brain was attributed to ∼24 days (Menzies and Gold, 1971). Striatum, which considered as one of the metabolically active regions of the brain (Brown et al., 2002), the exact half-life of mitochondria remains unknown. Although Rhes appears to regulate basal mitophagy, but the functional relevance of this yet to be determined. Rhes KO mice are hyperactive to dopaminergic drugs, but whether the lack of mitophagy may contribute to such hyperactive phenotype largely unknown. What is clear is Rhes is necessary for 3-NP induced mitophagy upregulation in the striatum. It is long known that 3-NP elicits striatal-specific lesion (Beal et al., 1993; Cirillo et al., 2019), and mechanisms such as oxidative stress and excitotoxicity were implicated (Albin and Greenamyre, 1992; Beal, 2000). Oxidative stress or excitotoxity can also occur in the cortex, however, 3-NP does not elicit lesion in the cortex (Fig. S5). Therefore, the molecular details that contributes to 3-NP induced striatal lesion was remain enigmatic. Now this study, provides a clear molecular route for striatal lesion by 3-NP. Our study defines the Rhes-mitophagy pathway as a principle trigger for the neuronal loss induced by mitochondrial toxin in the striatum. Our data also suggest that the major role for Rhes is to protect neurons by removing damaged mitochondria via mitophagy. However, upon exposure to irreversible toxins, such as 3-NP, which irreversibly damages the mitochondria, the Rhes-mediated mitophagy process are exacerbated and lead to depletion of the mitochondria and neuronal death (Fig. 10).

**Figure. 10.**
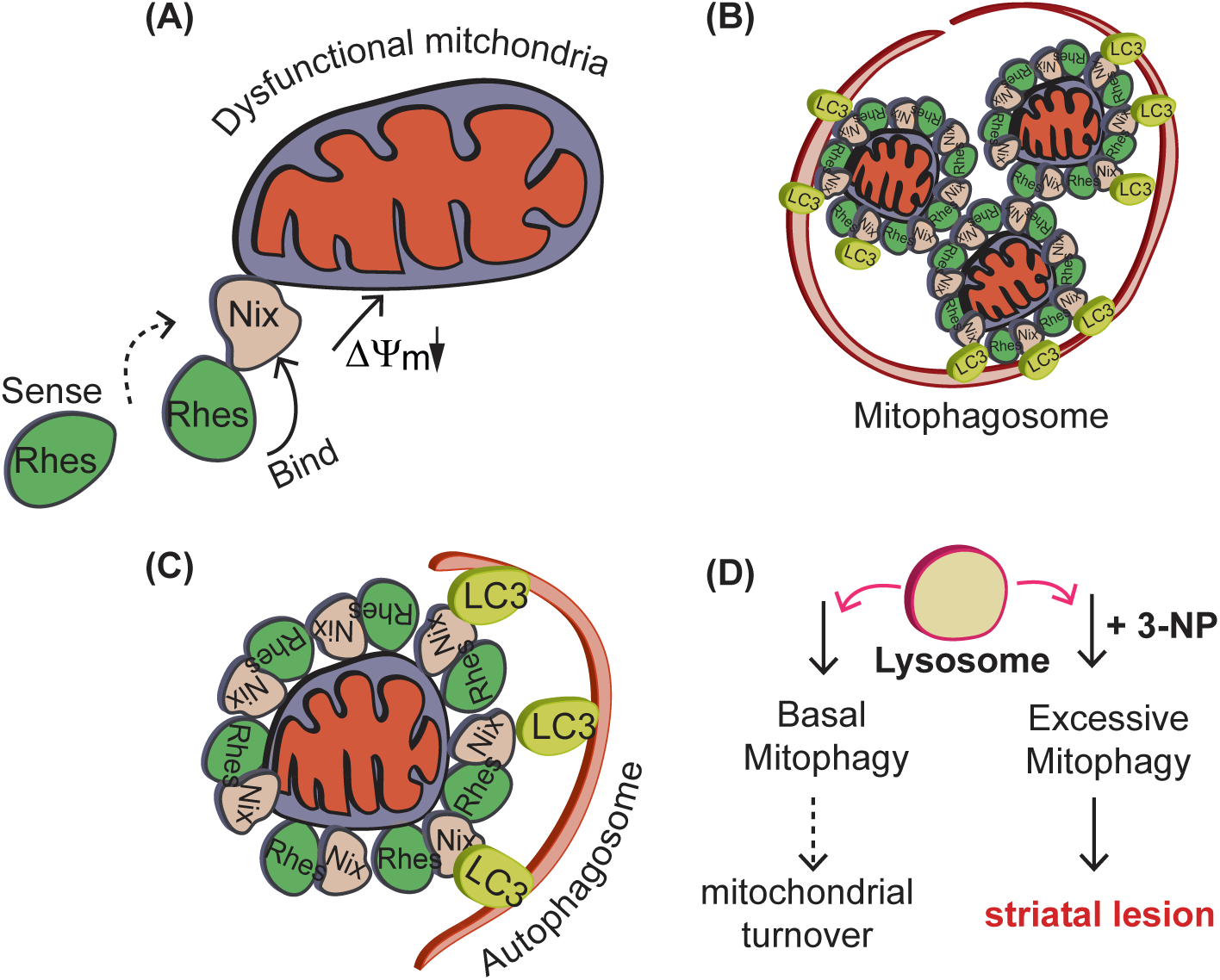
Model for Rhes-mediated mitophagy in the striatum. (A) Rhes sense dysfunctional mitochondria, interacts with Nix and diminishes membrane potentials. (B) Rhes-Nix interactions recruits autophagosomes via LC3. (C) Formation of mitophagosomes. (D) Rhes promotes mitochondrial degradation. Basal mitophagy may lead to mitochondrial turnover, excessive mitophagy in presence of mitochondrial toxin, 3-NP, promote striatal lesion.

Based on the data presented in this report, our current model predicts that Rhes may sense the dysfunctional mitochondria and binds to Nix to disrupt the ΔΨ*_m_* and initiate mitophagy. Rhes and Nix together may alter the membrane permeability pore leading to the diminishment of ΔΨ*_m_*. Nix alone cannot disrupt ΔΨ*_m_*, because there was no difference in TMRM signal intensity between Nix-WT and Nix-KO cells with or without 3-NP (Fig 8). Moreover, as mentioned above, Nix alone cannot elicit mitophagy in vivo because the levels of Nix between WT and Rhes KO striatum was similar, and yet KO show diminished mitophagy compared to WT (Fig. S6B). Mechanistically, as loss of ΔΨ*_m_* is a major initial step in the mitophagy upregulation, we hypothesize that altering the ΔΨ*_m_* by Rhes is the prominent step to initiate mitophagy. Rhes’ effect on mitophagy appears to be specific to 3-NP, but not FCCP (Fig 2) or rotenone (data not shown). How does this specificity is accomplished? One possibility would be that that Rhes, via yet unknown mechanisms, may particularly react to dysfunctional complex-II of mitochondria. Analogous to this, some factors in the substantia nigra may sense MPTP- or rotenone- induced mitochondrial dysfunction related to complex-I. Such tissue-specific regulator may set the stage for eliciting tissue-specific vulnerability through mitophagy. But the identity of such regulators in the substantia nigra remains unknown.

The implication that Rhes can travel from cell-to-cell and interacts with damaged mitochondria raises some intriguing possibility that Rhes may act like “mitochondrial surveillance factor.” We imagine that Rhes “scans” for damaged mitochondria in the surrounding cells and travel there via TNT-like membranous tubes to eliminates them. Conceptually such process in brain sounds like a “science fiction.” Considering the complexity of brain, the “surveillance” ability of Rhes may be needed for an effective co-ordination of billions of densely packed neurons in the brain.

Collectively, this study, reveals Rhes as a novel mediator of mitophagy and its impact on the regulation of striatal vulnerability. Development of therapeutic approaches that modulates mitophagy in the striatum may offer therapeutic opportunities for dysfunction related to striatal region of the brain.

## Acknowledgements

Dr. Long Yan of Max Plank Institute of Neuroscience, Jupiter, Florida, for imaging help; Alta Johnson and Bivian Torres of Flow cytometry core, Scripps Research, Jupiter, Florida, for helping in cell sorting and data analysis. We thank Sumitha Rajendra Rao and Stephen Zorc for technical assistance. The technical assistance of Rodolfo Paredes in the electron microscopic procedures is acknowledged. This part of the work was supported by Dirección General de Asuntos del Personal Académico, UNAM (project IN206719). A Training Grant partially supported this research in Alzheimer’s Drug Discovery from the Lottie French Lewis Fund of the Community Foundation for Palm Beach and Martin Counties. This research was supported by funding from NIH/NINDS R01-NS087019-01A1, NIH/NINDS R01-NS094577-01A1 and Cure for Huntington Disease Research (CHDI).

## Author contributions

M.S made initial observations. S.S further conceptualized and co-designed the project with M.S. M.S carried out all the experiments related to cell culture, primary neuron culture and interaction studies. U.Z carried out *in vivo* experiments such as 3-NP injections, behavior study and electron microscopy in collaboration with R.T. O.R assisted in interaction studies. M.K helped in seahorse experiment execution and data interpretation. M.E contributed efforts in sucrose density gradients. N.S did mouse work and analyzed Nix level in mice brain. V.S assisted in western blotting and DNA preparation. M.S, U.Z and S.S analyzed the data. S.S wrote the manuscript with the input from M.S and U.Z.

## Materials and methods

### Cell culture and chemicals

Mouse normal striatal neuronal cells (STHdh^Q7/Q7^) (Trettel et al., 2000) were cultured in growth medium containing Dulbecco’s modified Eagle’s medium (Thermo Fisher Scientific) with 10% fetal bovine serum (FBS), 1% penicillin-streptomycin, as described in our previous works (Pryor et al., 2014; Shahani et al., 2016; Subramaniam et al., 2009).

### Primary neuron culture

Primary neuron culture was prepared as described before (Sharma et al., 2019). In brief, animals were cared in accordance to the guidelines set forth by the National Institutes of Health regarding the proper treatment and use of laboratory animals and with the approval of Institutional Animal Care and Use Committee of The Scripps Research Institute. Striata of postnatal C57BL16 mice (P1) were removed and digested at 37°C for 15 min in a final concentration of 0.25% papain and resuspended in neuronal plating media (Neurobasal-A media, Thermo Fisher Scientific), with 5% FBS, 0.5 mM glutamax and 1% penicillin-streptomycin. Tissues were dissociated by trituration with a pipette. Further, cells plated in 35 mm glass bottom dishes (Matsunami D11140H) coated with 100 µg/ml poly-D lysine at the density of 2X105 cells per dish. Dishes were maintained in a 37°C, 5% CO2 incubator. After the cells adhered (1–3 h after plating), plating media was replaced with growth media (Neurobasal-A media, 2% B27, 0.5 mM glutamax and 1% penicillin-streptomycin).

### Antibodies, chemicals and treatments of cells

Lamp1 antibody was purchased from DSHB (AB 528127). GFP (sc-9996), VDAC1 (sc-390996), Myc (sc-40), GST (sc-138 HRP) and Actin monoclonal antibody (sc-47778) were obtained from Santacruz Biotechnology. mTOR (2983), SDHA (11998), LC3B (3868), Nix (12396) and Bip (3177) antibodies were from Cell Signaling Technologies. Rasd2 (Rhes) antibody (RHES-101AP) was obtained from FabGennix. mCherry antibody (NBP2-25157) was purchased from Novus Biologicals. Alexa 568 anti-mouse antibody was purchased from Thermo Fisher Scientific. Cyclohexamide (CHX) (7698), Chloroquine (C6628), Carbonyl cyanide 4- (trifluoromethoxy) phenylhydrazone (FCCP) (C2920) and 3-NP (N5636) were purchased from Sigma-Aldrich. CHX was dissolved in 95% ethanol and used at 100 µM for indicated time point. To assess autophagy flux, cells were pretreated with chloroquine (50 µm) for 4 hr, then proceeded for either vehicle or 3-NP treatment. 3-NP was dissolved in water, pH was adjusted to 7.4 and used at 10 mM for indicated time point. To assess mitophagy process in CHX chase experiment, cells were transfected with GFP or GFP-Rhes for 48 hr. Cells were treated with CHX 100 µM for indicated time point. Vehicle or 3-NP (10 mM) was added to culture for 1 hr and samples were harvested for western blotting. FCCP was dissolved in DMSO and used at 1 µm concentration for 6 hr. For virus infection, Striatal neuronal cells were infected with either adenovirus-CMV-null (abmgood) or adenovirus-CMV-Rhes (1 × 10^10^ opu/μl) at 1 MOI for 32-36 hr in complete media. Mitotracker (M7511) and Lysotracker (L12492) were obtained from Thermo Fisher Scientific. For mitochondrial staining, mitotracker was dissolved in DMSO and used at 200 nM. Lysotracker was used at 100 nM concentration. Mitotracker and/or Lysotracker was added for 30 mins after 48 h of GFP or GFP-Rhes transfection; cells were washed with D-PBS and live cell imaging was performed using live cell imaging solution (Thermo Scientific). TMRM (M20036) and propidium iodide (P3566) were obtained from Thermo Scientific and used as per manufacturer’s recommendation.

### Mitochondrial respiration analysis (Seahorse assay)

Striatal neuronal cells were plated in Sea Horse V7 culture plates and infected with Ad-null or Ad-Rhes (1 MOI for 32 hr) treated with vehicle or 3-NP (10 mM for 2 hr). To assess mitochondrial respiratory function, the growth media was replaced with XF media (unbuffered DMEM, pH 7.4, Agilent Technologies, Santa Clara, CA, USA; catalog # 103334-100) with added 10mM pyruvate and 2.5mM glucose. The plate could equilibrate for 30 minutes at 37^°^C in a CO_2_-free incubator before being loaded in the XF-96 analyzer according to the manufacturer’s instructions. Once in the analyzer, the plate was combined with the pre-calibrated XF cartridge containing the individual well ports A, B and C loaded with 10x stocks of the following chemicals, respectively: A. Oligomycin (20µg/ml); B. FCCP (20µM); C. Rotenone and Antimycin (20µM each). Oxygen consumption rate was monitored in real-time at 37^°^C under basal conditions and following the sequential delivery of the compounds in the cartridge ports, achieving their final 1x working concentrations. Non-mitochondrial rotenone/antimycin-insensitive OCR was subtracted from all other OCR measurements.

### Plasmids and Transfection

For GFP-Rhes and GFP-Rhes C263S, we amplified their respective cDNA from pCMV-Myc-Rhes (Subramaniam et al., 2009) and cloned it in EGFP-C1 vector. For GST-Rhes and GST-Rhes domains or mutants, their respective cDNAs were amplified from pCMV-Myc-Rhes (Subramaniam et al., 2009) and cloned in pCMV-GST vector. The following plasmids were obtained from Addgene (Table 1). Striatal neuronal cells seeded in 35 mm glass bottom dishes or other plates were transfected 24 h later with cDNA constructs using lipofectamine 2000 (Invitrogen) in the ratio of 1:2 (DNA: Lipofectamine) and for Primary neurons, we used 1:3 (DNA: Lipofectamine). For 10 cm dish, we transfected ∼8 µg of DNA, and for 35 mm dishes ranging from 1 to 2 µg total DNA.

**Table-1.**

| Plasmids | Gift from | Addgene ID |
| --- | --- | --- |
| mCherry-TOMM20-N-10 | Michael Davidson, The Florida State University, Florida | 55146 |
| pBMN-mCherry-Parkin | Richard J. Youle, NINDS, Maryland | 59419 |
| Myc-Nix | Joseph W. Gordon, University of Manitoba, Canada | 100795 |
| pCMVTNT PINK1 C-myc | Mark R Cookson, National Institute on Aging, Maryland | 13314 |
| <u>mCh-Drp1</u> | Gia K. Voeltz, University of Colorado Boulder, Colorado | 49152 |
| Myc-Nix-CytoB5 | Joseph W. Gordon, University of Manitoba, Canada | 100756 |
| Myc-Nix-MaoB | Joseph W. Gordon, University of Manitoba, Canada | 100757 |
| pDEST-CMV GFP-LC3B | Robin Ketteler, Medical research council, United Kingdom | 123114 |

### Flow cytometry

For TMRM staining, we seeded striatal neuronal cells in 6 well plate. Next day cells were transfected with 2 µg of DNA (GFP or GFP-Rhes) or infected with Ad-null or Ad-Rhes at 1 MOI for 48 hr. Cells were treated with vehicle or 3-NP for indicated time point. Cells were washed with PBS and trypsinized, centrifuged at 1800 RPM and resuspended in 1XPBS. The TMRM dye was added at 1 µl/ml for 30 min, incubated on dancing rocker and later washed 3 times with 1XPBS, filtered through a 40-µm nylon filter and resuspended in flow buffer [1x PBS (Ca/Mg++ free), 1 mM EDTA, 25mM HEPES pH 7.0, 2% FCS/FBS (Heat-Inactivated), 10 units/mL DNase]. Similar strategy was used to stain the cells with propidium iodide (1 µl/ml for 30 min). Cells were analyzed by flow cytometry (BD Biosciences LSR Fortessa cell analyzer). Each experiment was performed three times (in duplicate), and 10000-20000 cells were recorded for each sample. Data was compensated with single color controls and plotted using FlowJo software.

### Immunofluorescence

All the images stained for mitotracker and/or Lysotracker are acquired on live cells. For SDHA staining in GFP-Rhes transfected striatal neuronal cells following protocol was performed. At the indicated times after the post-transfection, cells were washed in D-PBS and fixed for 20 min in 4% PFA (Electron Microscopy Sciences). The cells were permeabilized with methanol and labeled with rabbit anti-SDHA antibody; (1∶100, for 18 hours at 4°C). For Myc-Rhes (1:100, for 18 hr at 4°C) staining, we used similar protocol as described above. The Alexa Fluor® 568 secondary antibodies were purchased from Thermo Fisher Scientific and used at 1:500 for 1 hour at room temperature.

### Image processing and colocalization coefficient quantification

All the fluorescent confocal images were taken in Zeiss 880 microscope using 20X or 63X oil immersion Plan-apochromat objective (1.4 NA). Excitation was via a 405 or 561 or 633 nm diode-pumped solid-state laser and the 488 nm line of an argon ion laser. Time-lapse acquisitions were performed using a 63X oil-immersion lens (1.4 NA). Images of striatal neuronal cells used for 2.5D or 3D reconstruction were acquired with an optimal Z-step of 0.27 µm covering the whole cellular volume. colocalization Processing and analysis was performed with Zen software black/blue edition 2012. Three-dimensional analyses and remodeling were done by Zen 2012 black edition software. Rhes mediated regulation of mitophagy in neighboring cell was performed as following. Striatal neuronal cells were transfected with GFP-Rhes and sorted through flowcytometry sorter to enrich GFP-Rhes expressing cell population. These sorted cells were co-cultured with striatal neuronal cells (control or Nix depleted) which were pretreated with vehicle or 3-NP (10 mM for 6 hr). Before coculture, vehicle or 3-NP added media was exchanged with fresh media so that complex II inhibition could be restricted to Rhes negative cells (pretreated cells). After 18 hr of co-culture, cells were stained for mitotracker orange for 30 min in serum free media. After that Media was changed to live cell imaging solution and images were acquired in confocal microscope. GFP-Rhes puncta, migrated to neighboring cells were selected and their colocalization with mitotracker was analyzed in Zen software.

### Generation of Nix Knockout cell line

Nix KO cell line was generated using Nix CRISPR/Cas9 plasmids from Santa Cruz Biotechnologies. First, we transfected the striatal neuronal cells with Nix CRISPR/Cas9 plasmid (SC-419357) or CRISPR/Cas9 control plasmid (SC-418922) in the 10cm dish. After 48 h we sorted the cells based on GFP fluorescence and re-cultured them. We passaged them 2-3 time and prepared lysate to confirm the Nix protein depletion by western blotting using Nix antibody.

### Protein expression and Western blots

Mitophagy process was assessed by Western blotting, cells were seeded in 6 well plate and next day transfected with 2 µg of DNA (GFP or GFP-Rhes) or infected with Ad-null or Ad-Rhes at 1 MOI for 48 hr. Cells were treated with indicated drugs or chemicles and washed 3 times with PBS. Protein lysates were prepared and loaded 40 µg of protein on polyacrylamide gels. For protein-protein interaction experiments, GST or GST-Rhes or GST-Rhes fragments or Myc-Nix or Myc-Pink1 or mCherry Parkin or mCherry DRP1 (4 μg each) were transfected in HEK293 cells (10 cm dish), and, after 48 hr, cells were lysed in lysis/binding buffer (50 mM Tris, pH 8.0, 150 mM NaCl, 10%glycerol, 1.0% NP-40) with a protease inhibitor cocktail (Roche) and phosphatase inhibitor II (Sigma). Glutathione beads were added, and the lysates were then rotated in a cold room for at least 5 hours. The beads were washed 3 times in binding buffer without a protease inhibitor cocktail, the proteins were eluted with 2X LDS-loading buffer. Proteins were separated on 4-12% Bis-Tris Gel (Invitrogen), and then transferred to PVDF membranes and probed with the indicated antibodies. HRP-conjugated secondary antibodies (Jackson Immuno Research, Inc.) were probed to detect bound primary IgG with a chemiluminescence imager (Alpha Innotech) using enhanced chemiluminescence (ECL), from WesternBright Quantum (Advansta). Indicated band intensities were quantified with ImageJ software.

### Subcellular Fractionation

For subcellular fractionations, mice were euthanized and both striatum were dissected immediately and homogenized using a glass dounce homogenizer (5 loose and 5 tight strokes) in buffer A of mitochondria isolation buffer (Thermo Scientific 89874). and kept on ice for 2 minutes. Buffer C was added to each sample and mixed by inverting 5 times. The homogenates were centrifuged at low speed (700g) for 10 minutes to separate nuclei and tissue chunks. The supernatants were immediately loaded on top of 10-50% sucrose gradients and centrifuged at 40000 RPM (SW41Ti rotor) at 4 degrees for 2 hours. The gradients were fractionated manually (11 X 1 ml fractions). Using methanol/chloroform, total protein of each fraction was precipitated. The protein pellets were resuspended in 2X LDS buffer and used for western blotting experiments. All the samples were proceeded at the same time. During developing the blots, all the groups were developed at the same time with corresponding antibody. For example, all the groups probed for SDHA, were developed together at the same time to get equal exposure for further analysis.

### Mice

For in vivo experiments we used Rhes-knockout (RhesKO) mice and C57BL/6J mice as control group. RhesKO mice were obtained from A. Usiello (Errico, 2008) and were backcrossed with C57BL/6J mice at least eight generations, homozygous RhesKO we used for all the experiments. Wildtype mice were obtained from Jackson Laboratory and maintained in our animal facility according to IACUC instructions. 3-NP injections were based on previous studies (Mealer et al, 2013; Blum et al., 2003). Briefly, 3-NP was dissolved in sterile PBS (0.1 M, 10 mg/ml), and pH was adjusted to pH 7.4 using sodium hydroxide. Mice received intraperitoneal injections twice a day (60 mg/Kg doses), with 2 h between injections for three consecutive days. In this study we used female and male mice, all of them were between16-20 weeks of age.

### Behavior evaluation

#### Beam walk test

Before injections, the mice were trained to walk over a wood beam (0.6 cm x 100 cm), suspended ∼50 cm from the benchtop with a safety box at one end. The training protocol was 4 days: 1 day traversing half the distance (50 cm) of the thickest beam (1.2 cm x 100 cm), and 2 days traversing the entirety of the beam (0.7 x 100 cm), and 1 day traversing the skinniest beam (0.6 x 100 cm). Training sessions consisted of 10 min habituation in the safety box, followed immediately by 4 attempts to cross the bar, each separated by 2 min to allow the mice to rest. After training, the daily performance was measured after a 10 min habituation period with 4 crossing attempts per mouse, using the skinniest beam. Baseline performance was measured on day 0 before injections, and testing was performed each subsequent morning before the next injection for 3 days. Sessions were videotaped and reviewed later for time to cross the 100 cm beam, the number of foot slips below the bar (errors), and the number of extra-supports under the beam used for crossing.

#### Open Field

Total activity was measured using the open field test. Each mouse was placed in the center of each open-top box (50 x 50 cm), under bright light and recorded via ceiling-mounted video camera for 10 min. Locomotor activity was analyzed using Ethovision XT11.5 animal tracking software (Noldus), total activity was measured on day 0 before injections, and each subsequent morning before rotarod, beam walk, and the next injection for 3 days. Because total activity can be affected for the continuous exposition of the mice to the test, based on preliminary results, animals were trained for 2 days before injections.

#### Rotarod

Alteration in motor coordination was determinate using the accelerating rotarod in wild type and RhesKO mice. After placed the mice on the rotating rod (diameter of 5 cm), they were tested using the accelerated rotarod from 4 to 40 RPM (cut off time of 300 s). Each mouse was tested three times and the average of the latency to fall was used for group analysis. Before injections, all the mice were trained for three days. Rotarod evaluation was measured on day 0 before injections, and each subsequent morning before the next injections for 3 days.

### Transmission electron microscopy and histology

After finishing the experiments, mice were fixed at 2 and 3 days of 3-NP treatment. For fixation, animals were anesthetized with tribromoethanol (1:50) in sterilized physiologic solution (NaCl 0.9%), and perfused transcardially with 15 ml of ice-cold 0.9% saline, followed by 20 ml of ice-cold 4% paraformaldehyde - 2.5% glutaraldehyde in 0.1 M PB, pH7.4. The brain was removed, post-fixed in the same solution. Right hemispheres were used for transmission electron microscopy (TEM) and left hemispheres for hematoxylin-eosin staining. For staining, hemisphere was cryoprotected in sucrose gradients (up to 30%), and sagittal striatal sections (40 µm-thick) were obtained.

The right hemisphere was used to transmission electron microscopy, after perfusion the striatum area containing cortical tissue was delimited. Then the tissue was osmicated (1% osmium tetroxide in PBS), washed with PB, dehydrated and embedded in epoxyresin. Semithin sections (500 nm) were obtained in an ultramicrotome (Reichert-Jung) and stained with Toluidine Blue for light microscopic observation, to identify the area of interest. Ultrathin sections were obtained and stained with Reynolds mixture (2% lead citrate and 2% uranyl acetate) and observed in a Jeol1200EXII transmission electron microscope. Electron micrographs were obtained and analyzed using the GATAN software. Mitochondrial alterations were determinate using the circularity index and mitochondrial area, which were quantified using the ImageJ software from at least 200 mitochondria from 4-5 mice. Organelle identification and quantification was made from around 40 neurons from 4-5 mice per group.

### Statistical analysis

Unless otherwise noted all experiments were carried out in duplicates and repeated at least three times. The statistical comparison was carried out between groups using one-way ANOVA, Student’s-*t*-test, two-way ANOVA and significance values were set at p<0.05, using graph pad Prism7.

**Movie S1.** Striatal neuronal cells were transfected with GFP-Rhes (green) and stained for mitotracker orange (red) and lysotracker (blue). Movie shows are time-lapse video of green, red and blue and its magnified area from selected rectangle region. Arrows shows the GFP-Rhes enrichment to globular mitochondria (also positive for lysotracker staining) in time dependent fashion. A total of six Z-stacks (planes) were acquired and presented as maximum intensity projection. Total time of movie file is 38 minutes. Twenty-eight cycles were collected at a rate of 1 minute 24 seconds per cycle. Movie was rendered slow motion with iMovie software.

## Supplementary Figure Legends

**Fig. S1.**
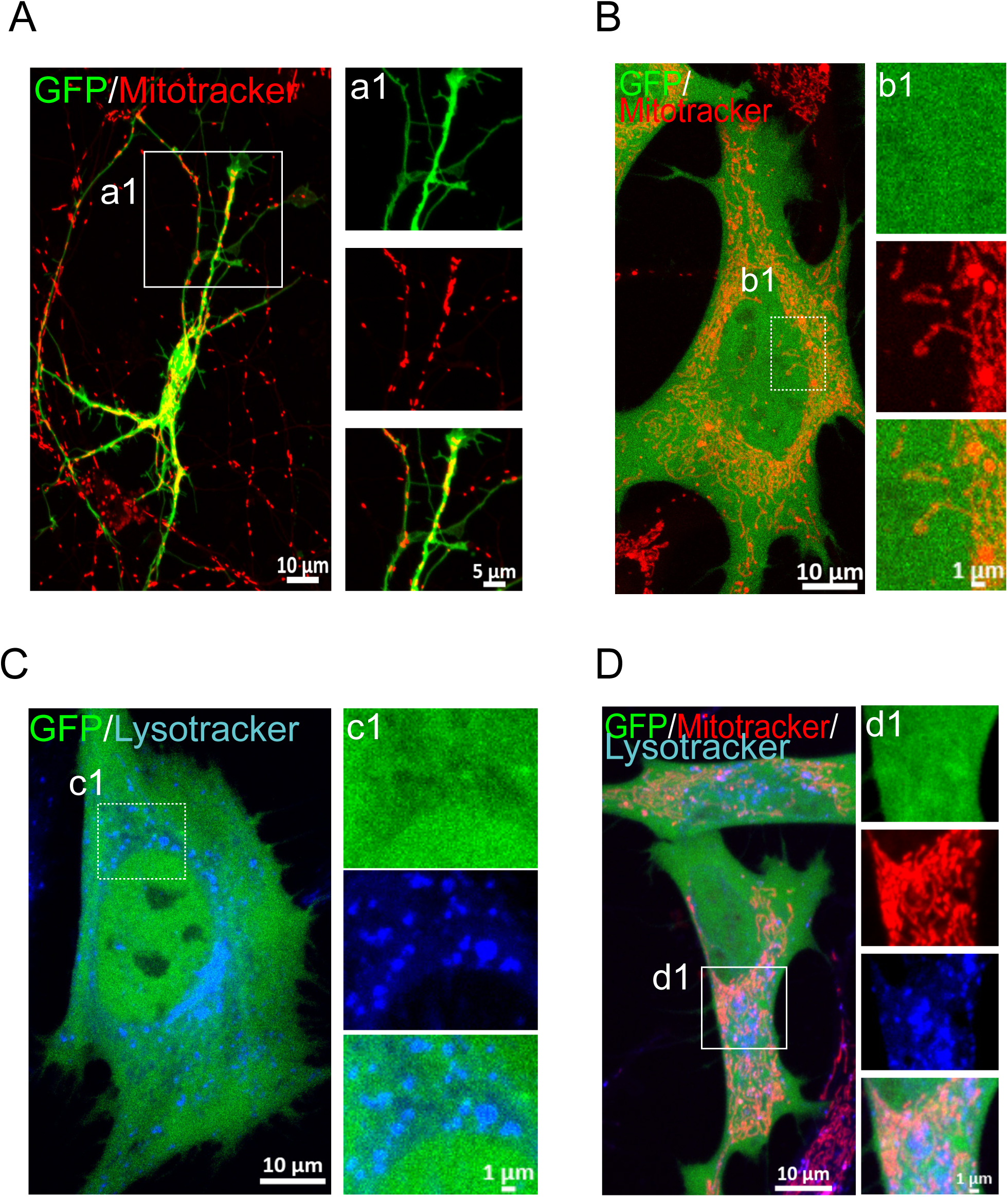
GFP control does not interact with mitochondria or lysosome. (A) Representative confocal image of primary striatal neuron transfected with GFP vector. Cells were stained with mitotracker orange (red). a1 is inset from selected region. (B) Confocal image of striatal neuronal cells transfected with GFP. Cells were stained with mitotracker orange (red) b1 is inset from selected region. (C) Confocal image of GFP transfected striatal neuronal cell. Cells were stained with lysotracker (blue). c1 is magnified area from selected region. (D) Confocal image of striatal neuronal cells transfected with GFP vector. Cells were stained for mitotracker orange (red) and lysotracker (blue). d1 is magnified area from selected region.

**Fig. S2.**
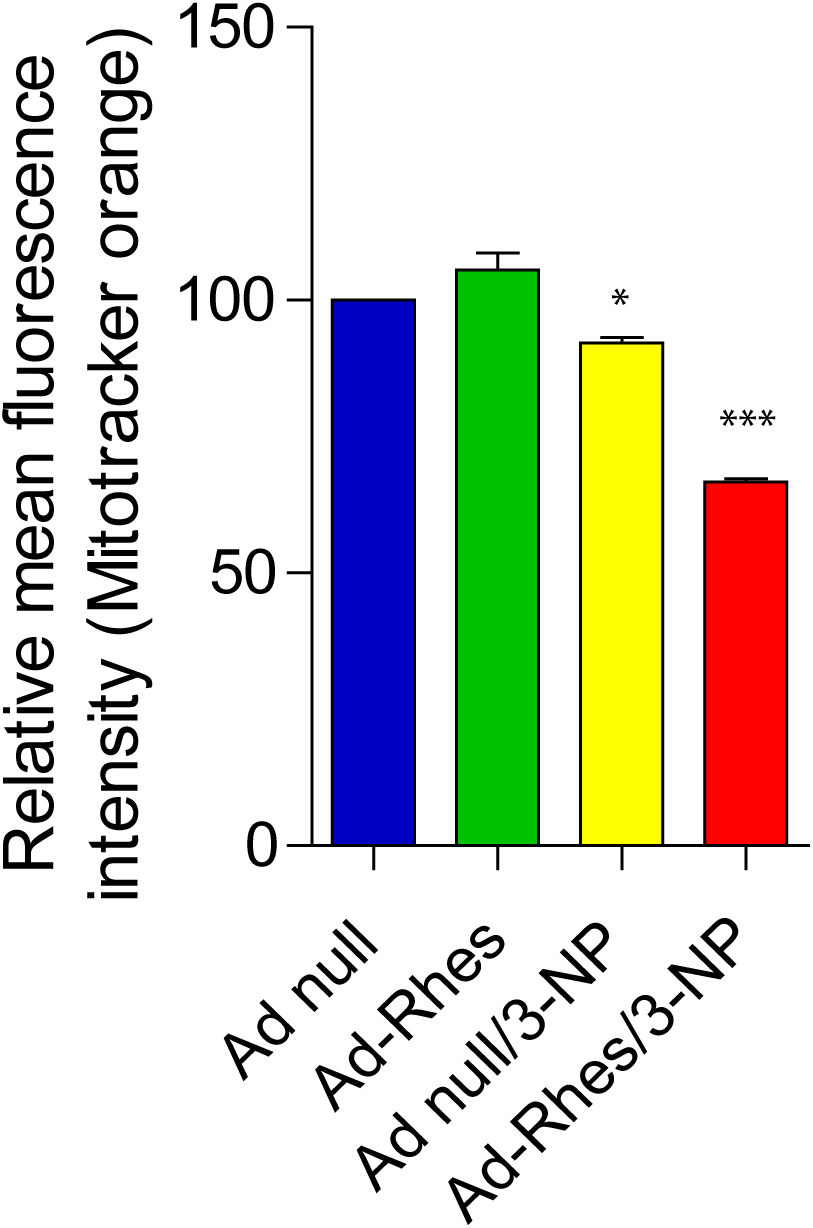
Rhes expressing cells show reduced mitotracker intensity in 3-NP treatment. Bar graph shows relative mean fluorescence intensity of mitotracker orange in striatal neuronal cells infected with Ad-null or Ad-Rhes, treated with vehicle or 3-NP (10 mM for 2 hr). Statistical comparison was done between groups using One-way anova test (compared to Ad-null/vehicle) and significance values were set at *p<0.05, ***p<0.001. Data mean ± SEM.

**Fig. S3.**
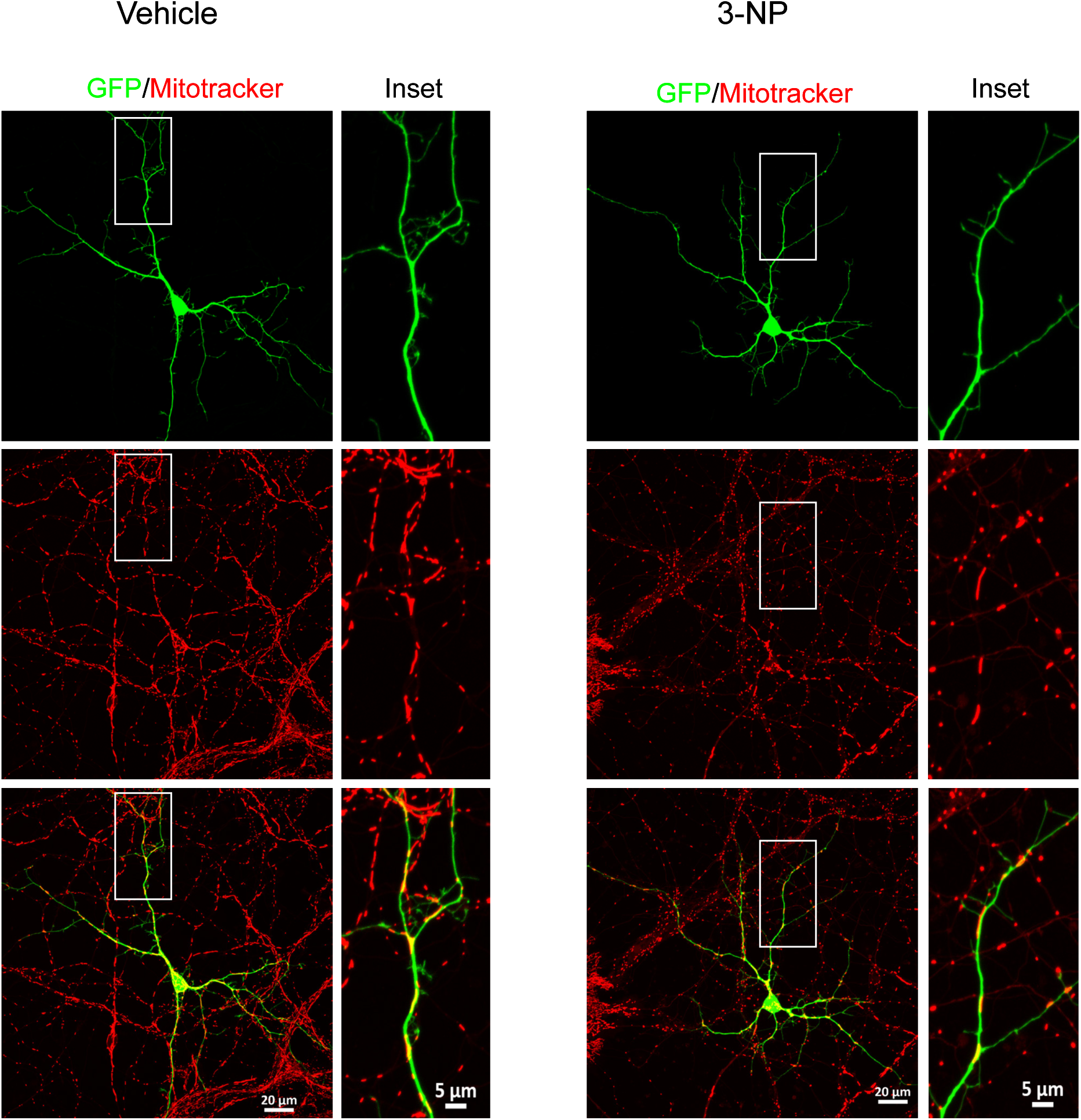
GFP alone doesn’t affect mitotracker staining in primary neuron, treated with 3-NP. Confocal image of primary striatal neuron transfected with GFP vector, treated with vehicle or 3-NP (10 mM and for 16 hr). Corresponding insets show the magnified area from selected region.

**Fig. S4.**
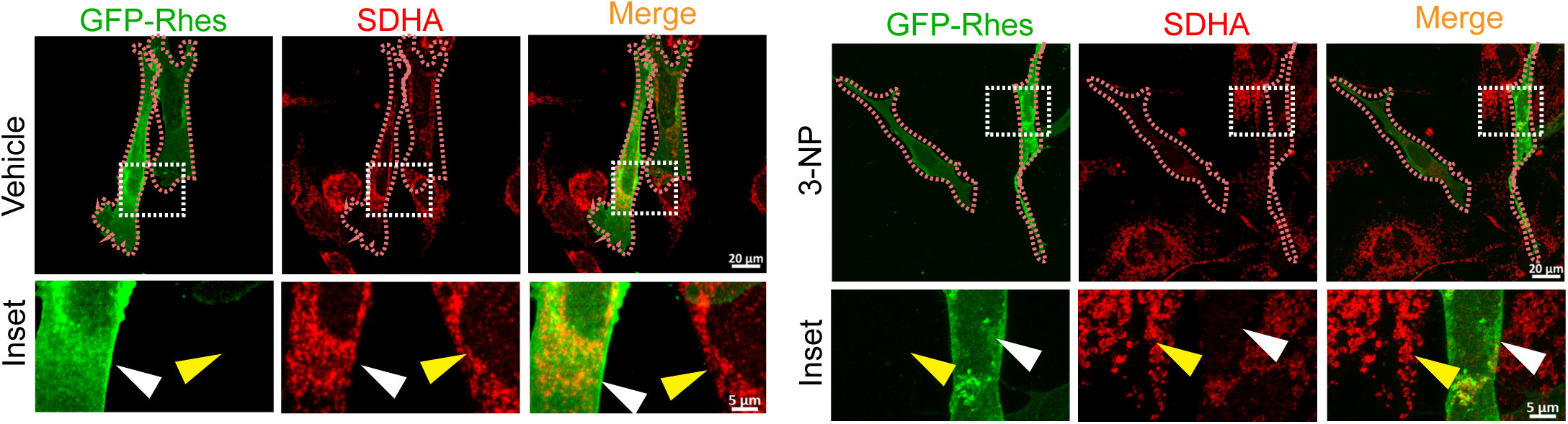
Rhes diminishes SDHA signal in presence of 3-NP. Representative confocal image of striatal neuronal cells transfected with GFP-Rhes, treated with vehicle or 3-NP and stained for endogenous SDHA (a mitochondrial protein). GFP-Rhes transfected cell periphery was marked by dashed line (brown). Insets show the magnified region from the selected area. White arrowhead represents the GFP-Rhes transfected cell and yellow arrowhead shows the untransfected cell in the same area.

**Fig. S5.**
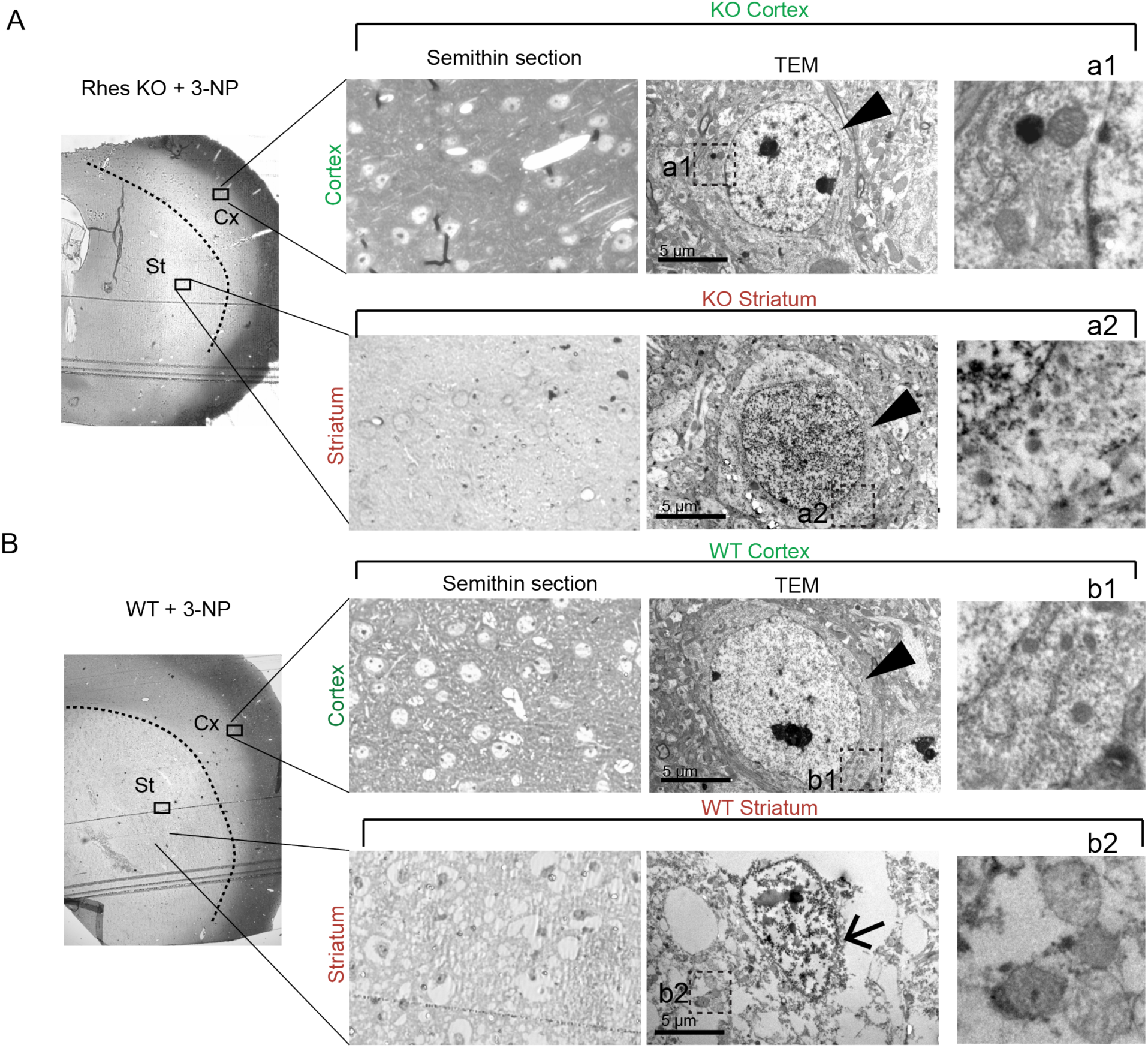
3-NP does not promote mitophagy or lesion in cortical neurons. (A, B) Representative electronic micrographs from striatal and cortical neurons after 3 days of 3NP treatment form Rhes KO (A) and WT brain (B). Arrowhead indicates intact neuronal morphology; arrow indicates loss of nuclear integrity and abnormal shrunken nuclei. Insets a1, a2, b1 and b2 indicates mitochondrial morphology in cortical and striatal areas of Rhes KO and WT mice brain.

**Fig. S6.**
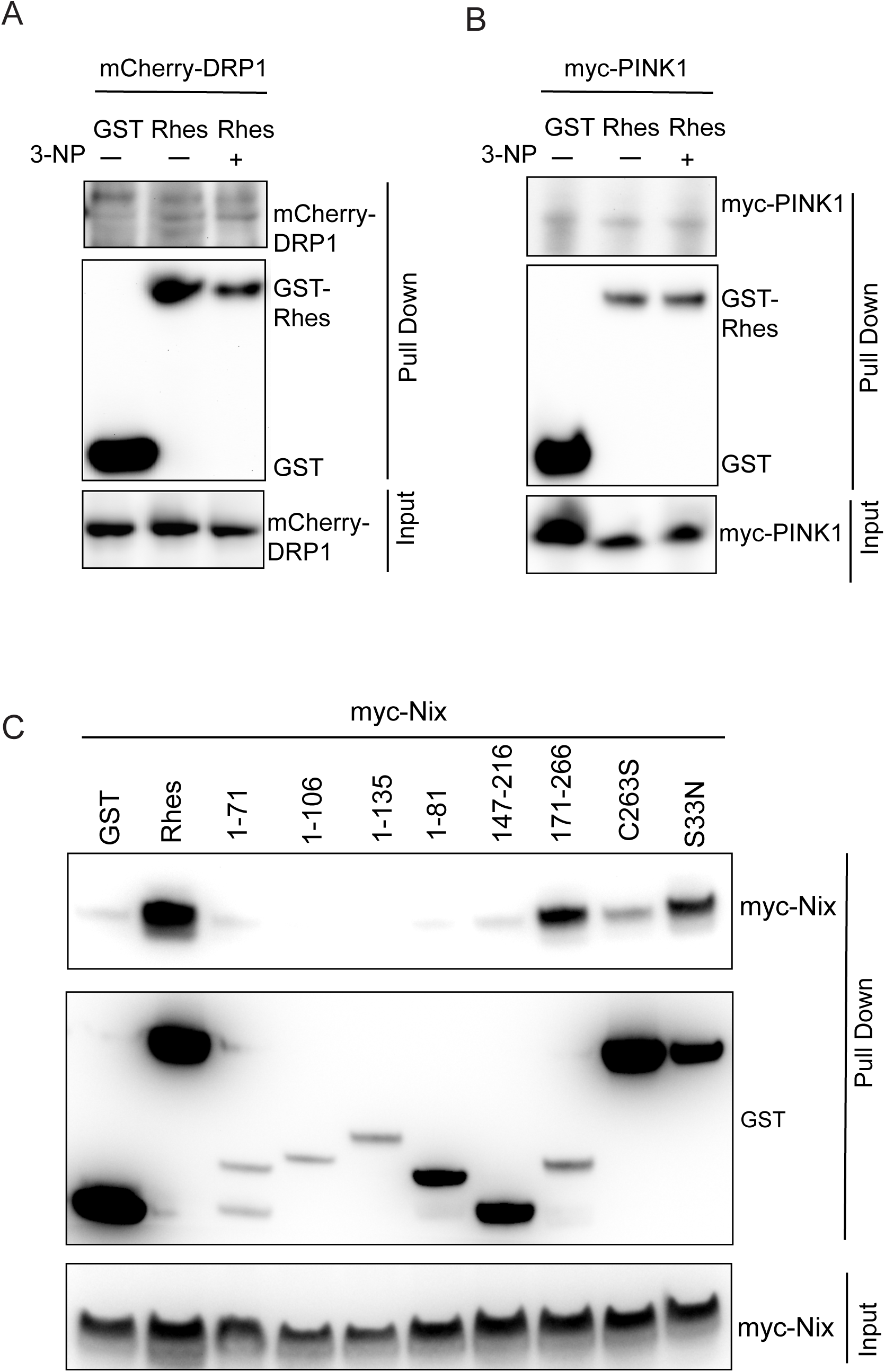
A. **Rhes does not interact with DRP1.** (A) HEK 293 cells were co-transfected with GST or GST-Rhes and mCherry-DRP1. Cells were treated with vehicle or 3-NP for 2 h, lysed and protein lysate was pull down using GST-sepharose beads and processed for immunoblotting. Blots were probed for mCherry and GST antibody to detect mCherry-DRP1 and GST or GST-Rhes in pulled down samples. Input was loaded as 5% of total lysate. B. **Rhes does not interact with PINK**1. HEK 293 cells were co-transfected with GST or GST-Rhes and myc-Pink1. Cells were treated with vehicle or 3-NP for 2 hr and processed as describe above. Blots were probed for myc and GST antibody to detect myc-Pink1 and GST or GST-Rhes in pulled down samples. Input was loaded as 5% of total lysate. C. **Rhes binds to Nix via SUMO E3 like and CAAX domain**. HEK 293 cells were co-transfected with myc-Nix and GST or GST-Rhes or its domain and mutants as indicated. After 48 hr of transfection cells were lysed and protein lysate was pull down using GST-sepharose beads and processed for immunoblotting. Blots were probed for myc and GST antibody to detect myc-Nix and GST or GST-Rhes or Rhes domain and mutants in pulled down samples. Input was loaded as 5% of total lysate.

**Fig. S7.**
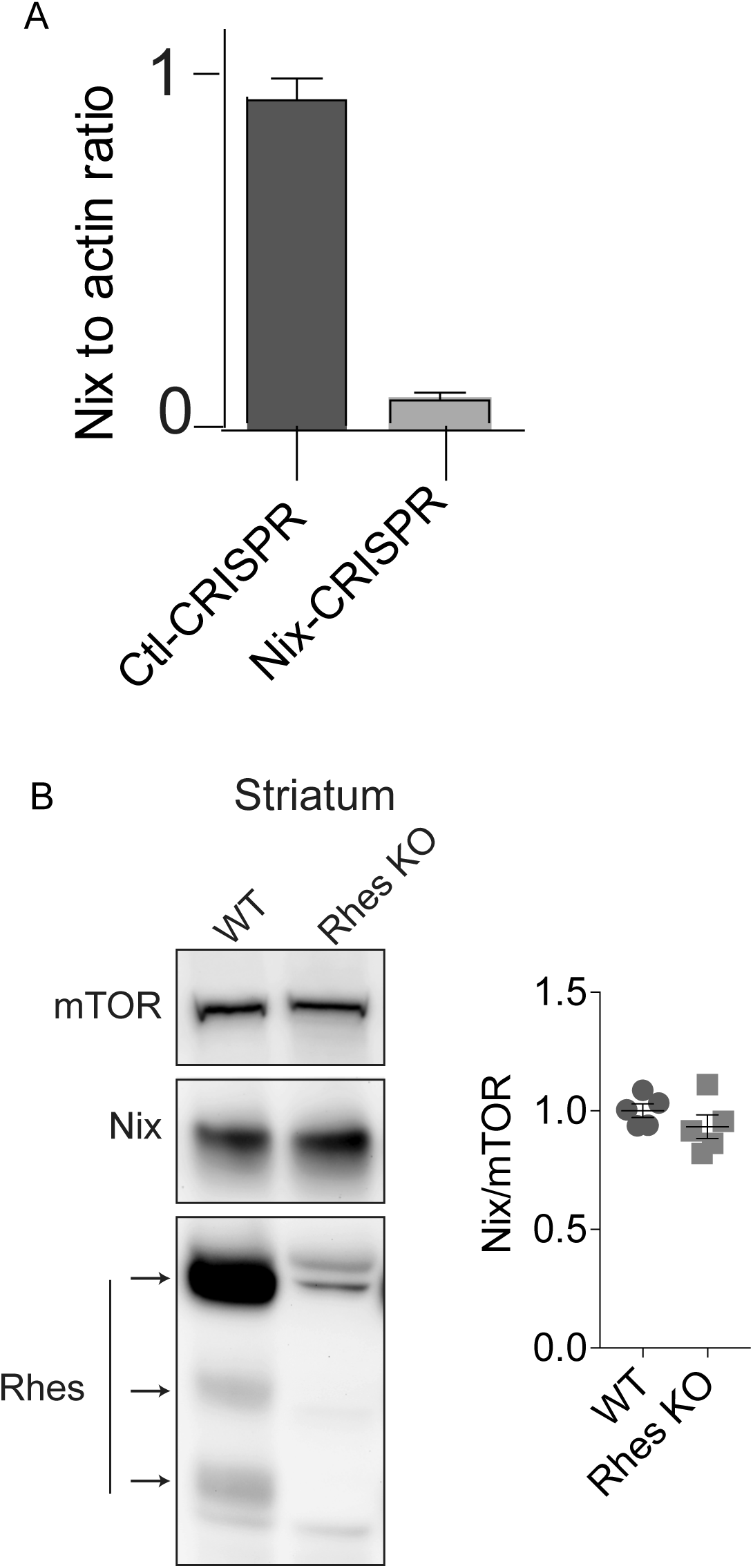
(A). **Efficient depletion using CRISPR/Cas-9 tool.** Bar graph shows the quantification for Nix protein normalized to Actin in CRISPR/Cas9 mediated control or Nix depleted striatal neuronal cells. ***p < 0.001 vs. control CRISPR. (B). **Nix levels are unaltered in the striatum of Rhes KO.** Protein lysates from Wt or Rhes KO mice striatum were probed for mTOR, Nix and Rhes protein. Bar graph shows the Nix protein levels (normalized to mTOR) in Wt or Rhes KO striatum (n=5 mice per group).

